# Voltage-sensor trapping of cardiac Na^+^ channels by Mg-protoporphyrin impairs cancer cell migration

**DOI:** 10.1101/2025.07.28.667138

**Authors:** Mahdi Jamili, Marwa Ahmed, Alisa Bernert, Johann Rößler, Guido Gessner, Roland Schönherr, Toshinori Hoshi, Stefan H. Heinemann

## Abstract

The human voltage-gated sodium channel hNa_V_1.5 is essential for cardiac excitability. Though underrecognized, Na_V_1.5 is also expressed in multiple cancers, promoting cell migration and malignancy. hNa_V_1.5 is a therapeutic target but limited isoform specificity presents a risk of side effects via neuronal and skeletal muscle Na_V_ channels. Here we identify Mg^2+^-protoporphyrin IX (MgPpIX), a Mg-containing tetrapyrrole and intermediate in chlorophyll biosynthesis, as inhibitor of hNa_V_1.5 (IC_50_ of 1 nM). The activity profile of various metal protoporphyrins correlates with the electrostatic potential at the metal center of the compounds. MgPpIX is specific to hNa_V_1.5, as no inhibition of other hNa_V_ isoforms (hNa_V_1.2, 1.4, 1.7, 1.8) was detected. A mutagenesis study and structural modeling reveals that MgPpIX stabilizes the domain-II voltage sensor in the deactivated conformation, with residues E795 and N803 being relevant determinants. MgPpIX also inhibits native Na_V_ channels in breast cancer MDA-MB-231 and colorectal carcinoma SW480 cell lines, and suppresses cell migration. MgPpIX is an exceptionally potent and specific inhibitor of hNa_V_1.5 and may serve as a lead compound in anti-cancer drug development.

## Introduction

Voltage-gated sodium channels (Na_V_ channels) are crucial in cells with action potentials, such as neurons, muscle cells, and some neurosecretory cells. Nine genes in the human genome code for pore-forming α subunits (Na_V_1.1-1.9, genes *SCN1A-SCN11A*). Na_V_1.5 (*SCN5A*) is primarily found in the heart, while Na_V_1.4 is present in skeletal muscle, Na_V_1.1 and Na_V_1.2 in the central nervous system, and Na_V_1.7-Na_V_1.9 in the peripheral nervous system. Na_V_ gain-of-function gives rise to a plethora of hereditary and acquired diseases, such as myotonia, epilepsy, cardiac arrhythmia, and erythromelalgia (Savio-Galimberti *et al*, 2012). Loss of Na_V_ function can also have severe consequences, as exemplified in various forms of epilepsy (Na_V_1.2) (Menezes *et al*, 2020), indifference to pain (Na_V_1.7) (Lischka *et al*, 2022), and Brugada syndrome (Na_V_1.5) – a form of cardiac arrhythmia (Li *et al*, 2020).

In basic research and in human medicine, molecules that specifically affect Na_V_ isoforms are of utmost importance (Daimi *et al*, 2022; de Lera Ruiz & Kraus, 2015). The potent Na_V_ channel blocker tetrodotoxin (TTX) has proven useful in discriminating among TTX-sensitive (Na_V_ isoforms 1-4, 6, 7), less sensitive (Na_V_1.5), and insensitive (Na_V_1.8, Na_V_1.9) channels. Most pharmacological agents targeting Na_V_ channels aim to decrease the electrical excitability, slowing action potential firing in neurons or in cardiac cells, or shortening action potential duration by blocking the so-called late Na^+^ currents. Further, local anesthetics and antiepileptics diminish Na_V_ channel activity in a use-dependent manner (Körner *et al*, 2022). The recent approval by the US Food and Drug Administration of suzetrigine, a small-molecule pain medication specifically targeting Na_V_1.8, illustrates the clinical importance of developing isoform-specific Na_V_ modulators (Osteen *et al*, 2025).

Na_V_ channels are predominantly expressed in excitable cells; however, their presence in non-excitable cell types, including Schwann cells (Shrager *et al*, 1985), microglia (Black *et al*, 2009), macrophages (Carrithers *et al*, 2007), and various cancer cells indicates a role for Na_V_ channels in non-excitable tissues (Brackenbury, 2012; Liu *et al*, 2024). Upregulation of Na_V_ α and β subunits has been reported in multiple cancer types, including those derived from breast, colon, lung, prostate, and ovarian tissues as well as in lymphoma, and melanoma cells (Brackenbury, 2012; Fraser *et al*, 2005; Liu *et al*., 2024; Roger *et al*, 2015), with hNa_V_1.5 playing a prominent role (Rajaratinam *et al*, 2022).

Structurally, Na_V_ channels are membrane proteins composed of four homologous domains (I-IV), each comprising six transmembrane segments (S1-S6), with the pore loops between S5 and S6 determining the selectivity for Na^+^ ions (Heinemann *et al*, 1992) and the blocking efficacy of TTX (Terlau *et al*, 1991), and S1-S4 forming the voltage-sensor domains (VSDs). The atomic structure of hNa_V_1.5 channels has been determined through cryo-electron microscopy (Li *et al*, 2021). Structural data also give insight the inhibitory action of pore-blocking drugs (Lenaeus *et al*, 2023).

Recently, we showed that extracellularly applied hemin (Fe^3+^-protoporphyrin IX, Fe(III)PpIX, an amphipathic compound with a hydrophobic porphyrin macrocycle) inhibits hNa_V_1.5 channels (Gessner *et al*, 2022). Hemin does not act as a channel pore blocker but rather as a gating modifier that affects protein conformational changes leading to channel opening. While hemin affects hNa_V_1.5-mediated Na^+^ current with an apparent half-inhibitory constant (IC_50_) of about 100 nM, heme (Fe(II)PpIX), a prosthetic group with key functions in redox reactions of the respective hemo-proteins (Ponka, 1999; Tsiftsoglou *et al*, 2006), has no impact on hNa_V_1.5 channels (Gessner *et al*., 2022). Thus, and importantly, the hemin–hNa_V_1.5 interaction is notably distinct from previously reported cases of heme being a modulator of ion channel function where both heme and hemin are similarly effective (Tang *et al*, 2003).

The clear contrasting effects of hemin and heme on hNa_V_1.5 indicate the metal center as a key determinant of a metal protoporphyrin’s (MePpIX) interaction with the channel. We thus investigated whether other MePpIXs might also inhibit hNa_V_1.5 channels. We found that Mg^2+^-protoporphyrin IX (MgPpIX), a naturally occurring tetrapyrrole intermediate in the chlorophyll synthesis pathway (Mochizuki *et al*, 2008), is by far the most potent and specific inhibitory gating modifier of human Na_V_1.5 channels, with an IC_50_ of about 1 nM. MgPpIX stabilizes the voltage sensor of domain II in the deactivated state, which critically depends on E795 and N803 facing the extracellular side and the positive charge of the metal center in MgPpIX. The unprecedented subtype specificity and its potency to slow down the migration of *SCN5A*-expressing cancer cells highlight the pharmacological potential of MgPpIX as an anti-tumor lead compound.

## Results

### Select metal protoporphyrins inhibit hNa_V_1.5 channels

Cardiac voltage-gated sodium channels (Na_V_1.5) are specifically inhibited by Fe(III)PpIX (Gessner *et al*., 2022). Here we address how the inhibitory action depends on the central metal ion in the protoporphyrin scaffold (Fig. 1A *top right*). To this end, human Na_V_1.5 channels were expressed in HEK293T cells, Na^+^ currents were repetitively measured at -30 mV, and the impact of various MePpIXs before and 5 min after application to the extracellular side at 1 µM was compared. Experiments were performed under ambient (oxidizing) conditions; the probable redox state of the MePpIXs is indicated (Fig. 1A). As we reported earlier, hemin (1 µM) decreased the peak inward current size to ∼20%. In striking contrast, only ∼1% of the current remained with MgPpIX (Fig. 1A, *top*) – a near total loss of current. GaPpIX and ZnPpIX were also more potent than hemin, with about 10% current remaining. MnPpIX was slightly less potent than hemin, and 1 µM of CuPpIX, NiPpIX, and SnPpIX failed to affect the current. Limited solubility precluded examination at higher MePpIX concentrations. The rank order of hNa_V_1.5 inhibition by MePpIX is summarized in Fig. 1B with MgPpIX being by far the most potent: Mg(II) >> Ga(III) ≈ Zn(II) > Fe(III) > Mn(III) >> Cu(II), Ni(II), Sn(IV). Chlorophyll-A (1 µM), which consists of a modified MgPpIX ring with an added hydrocarbon chain, had no detectable effect (Fig. 1B, *right* Chl); the Mg^2+^ core in a macrocycle alone does not fully determine the inhibitory potency. Any influence of residual free Mg^2+^ in the MgPpIX samples can be excluded because the extracellular buffer already contained 2 mM MgCl_2_. The inhibitory rank order correlates well with their DFT-derived electronic surface potential, with more electropositive metal centers leading to greater inhibition, as illustrated for Mg(II), Zn(II), and Ni(II) (Fig. 1C). For more information on DFT calculations and predicted electronic surface potentials, refer to Fig. EV1.

**Figure 1.**
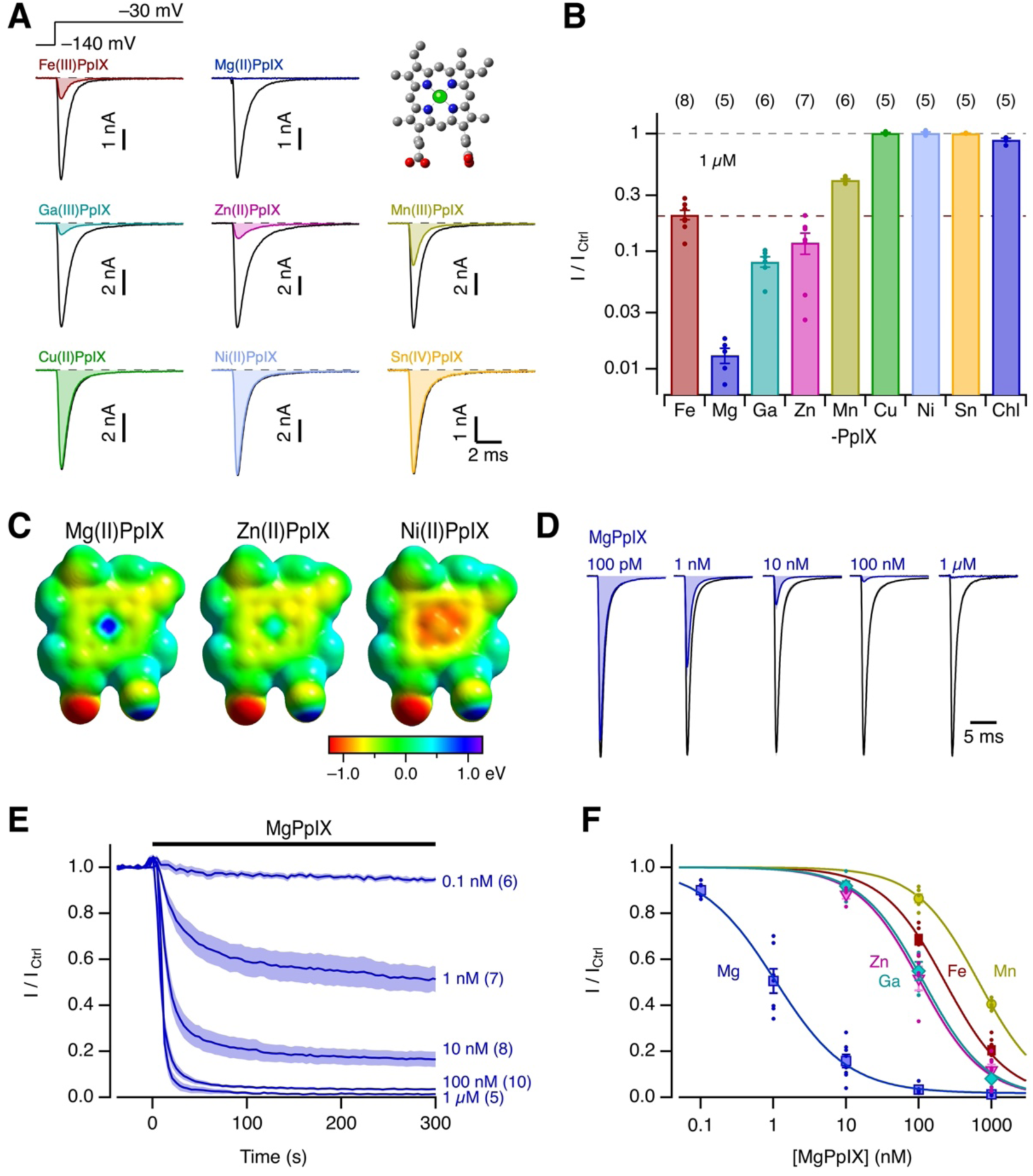
Inhibition of hNa_V_1.5 channels by metal protoporphyrins. (A) Whole-cell currents recorded using the indicated pulse protocol before (black) and 5 min after the application of 1 µM of the indicated metal protoporphyrins (MePpIX) (colored). The inset shows the structure of MePpIX, for which the central metal ion (green) was varied in this experiment. Mean remaining peak current after 5 min relative to the control in the presence of 1 µM of MePpIX. The last bar “Chl” shows the result for 1 µM chlorophyll-A. Bars indicate means ± sem with individual results indicated as dots. (**C**) Electrostatic surface potential of Mg(II)PpIX, Zn(II)PpIX, and Ni(II)PpIX according to DFT calculations using the aug-cc-pVTZ basis set; also see Fig EV1. (**D**) Mean current traces at -30 mV before (black) and after the application of the indicated concentrations of MgPpIX; data are normalized to the peak current of the control records (for *n* values, see (E)). (E) Time course of mean peak currents from records as shown in (D) with MgPpIX applied at time zero at the indicated concentrations. Mean values are shown as thick lines, sem is indicated by shading, and *n* is in parentheses. (**F**) Mean relative hNa_V_1.5 current after 5 min of MePpIX application as a function of concentration. The superimposed curves are fits to the Hill function (Eq. 4), for MgPpIX yielding an IC_50_ value of 1.05 ± 0.10 nM, a Hill coefficient of 0.86 ± 0.06, and a remaining fraction of 0.017 ± 0.015. Further IC_50_ values: ZnPpIX 106 ± 11 nM, GaPpIX 117 ± 8 nM, FePpIX 231 ± 18 nM, MnPpIX 672 ± 18 nM with the Hill coefficient constrained to 1 and the remaining current to zero.

MgPpIX caused a concentration-dependent reduction in hNa_V_1.5 current size, without affecting its time course (Fig. 1D). The onset of the inhibition also depended on concentration (Fig. 1E). Near full inhibition occurred within 10 s for 100 nM MgPpIX and higher, and even at 10 nM, a steady state was reached after about 200 s (Fig. 1E; for other MePpIX, see Suppl. Fig. 1). The concentration dependence of the relative remaining current after 5 min of MgPpIX application was fit with a Hill function (Eq. 4, Fig. 1F), yielding an apparent IC_50_ of 1.05 ± 0.10 nM, a Hill coefficient *n*_H_ = 0.86 ± 0.06, and a remaining fraction *a*_οο_ = 0.017 ± 0.015. For values of other MePpIX, see legend of Fig. 1.

MgPpIX might catalyze the production of reactive species, potentially targeting redox-sensitive residues in hNa_V_1.5. To rule out an influence of cysteine C373 in the pore region of domain I – a residue unique to Na_V_1.5 channels and responsible for the channel’s weak sensitivity to TTX – we examined variant hNa_V_1.5-C373Y. The current remaining after application of 100 nM MgPpIX was 5.4 ± 1.2% (*n* = 5), not markedly different from that of the wild type (3.4 ± 0.4%, *n* = 10, t-test *p* = 0.08; Fig. EV2).

### Mechanism of channel inhibition: reverse use-dependence

The inhibition of hNa_V_1.5 channels by hemin is characterized by a pronounced reverse use-dependence, with depolarizing pulses restoring channel activity (Gessner *et al*., 2022). To examine if MgPpIX shows a similar dependence, currents through hNa_V_1.5 were elicited by a family of depolarization pulses, producing voltage-dependent currents (Fig. 2A *left*). Under control conditions, preceding each depolarizing pulse with a 250-ms depolarizing prepulse to 40 mV did not affect the current (Fig. 2A *right*). With MgPpIX (100 nM), the currents at all voltages were abolished without the prepulse. With the prepulse, ∼50% of the control current was recovered at each voltage, indicating reverse use-dependence. The voltage and duration dependence of the prepulse required to recover hNa_V_1.5 currents in the presence of MgPpIX were quantified using prepulses of varying voltage (Fig. 2C) and duration (Fig. 2D). The half-maximal voltage (-21.8 ± 1.1 mV for 100 nM, -37.5 ± 1.5 mV for 10 nM) and the time constants for current recovery at 50 mV (52.7 ± 1.6 ms for 100 nM, 49.5 ± 1.9 ms for 10 nM MgPpIX) were only weakly dependent on the MgPpIX concentration. However, the kinetics for reestablishing the inhibition were accelerated by a factor of 4.3 with a 10-fold increase in MgPpIX concentration (Figs. 2E, EV3F). This concentration dependence is expected if the prepulse causes MgPpIX to dissociate from the channel binding site(s). Yet, hNa_V_1.5 inhibition was not readily reversed by removing MgPpIX from the bulk extracellular solution, even when depolarizing pulses were applied, suggesting a more complex mechanism, possibly due to MgPpIX ‘s hydrophobicity and thus partitioning in the lipid phase (Fig. 2F *top*). Washing with fatty-acid-free bovine serum albumin, a protein that provides hydrophobic binding pockets, resulted in partial recovery (Fig. 2F *bottom*).

**Figure 2.**
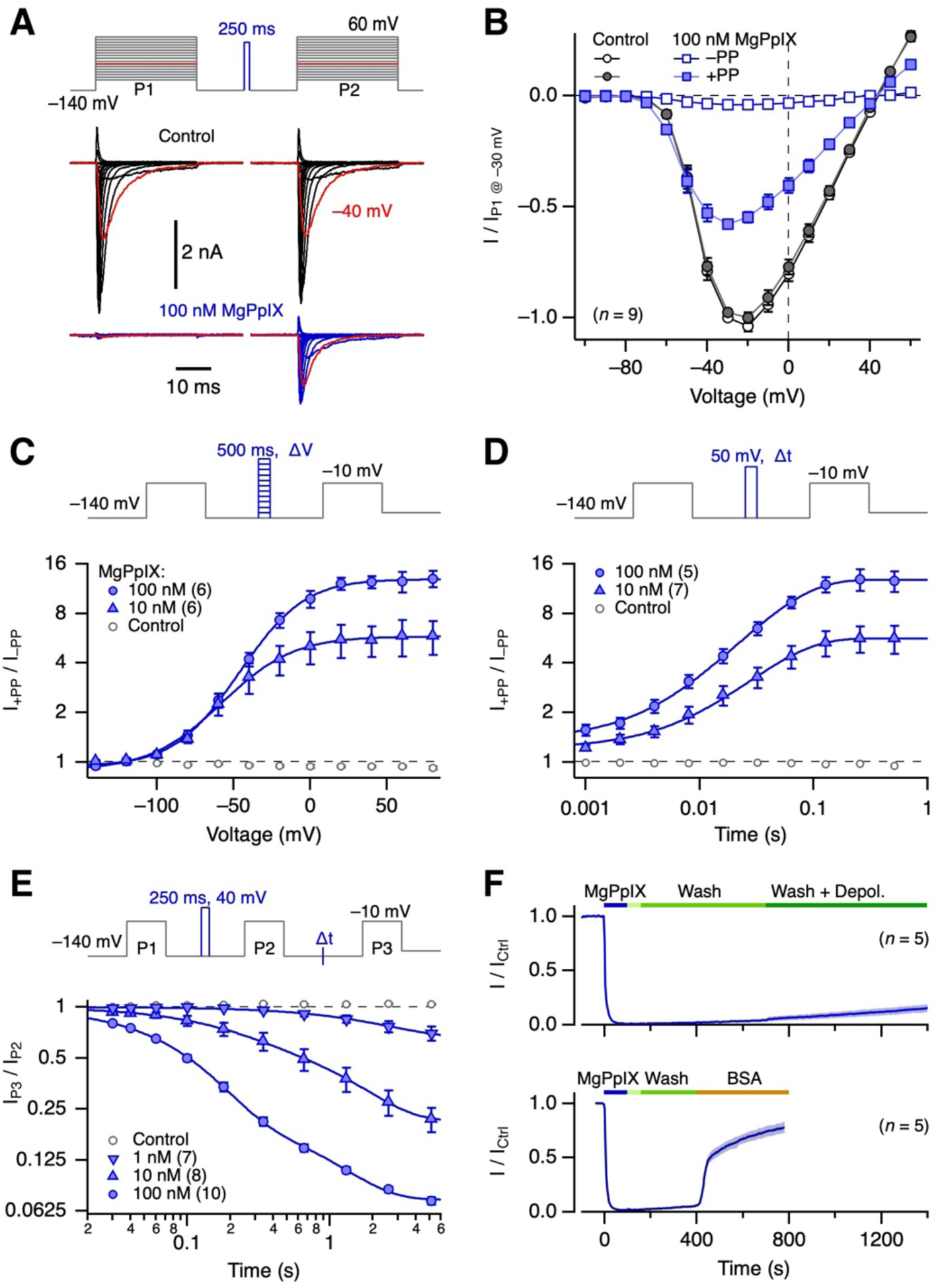
Reverse use-dependence of channel inhibition by MgPpIX. (**A**) Pulse protocol (*top*) and overlaid current traces from HEK293T cells expressing hNa_V_1.5 before (*bottom*, black) and after the application of 100 nM MgPpIX (blue). Between pulse 1 (P1) and pulse 2 (P2), the cell was depolarized to 40 mV for 250 ms. Traces with depolarizing pulses to -40 mV are shown in red. (**B**) Mean normalized peak currents from experiments as in (A) as a function of voltage, without (–PP) and with prepulse (+PP). Straight lines connect data points for clarity. (**C**) Pulse protocol (*top*) designed to evaluate the voltage dependence with which a 500-ms prepulse relieves the channel from inhibition. (*bottom*) Fractional recovery for peak current as a function of voltage, with superimposed Boltzmann fits (Eq. 3) for 10 and 100 nM MgPpIX. (**D**) Pulse protocol (*top*) for measuring the kinetics of recovery from MgPpIX-induced inhibition at 50 mV. (*bottom*) Fractional recovery of the peak current as a function of prepulse time at 50 mV, with superimposed single-exponential fits. (**E**) Pulse protocol (*top*) for measuring the kinetics of channel inhibition at -140 mV after prepulse-induced reversal. (*bottom*) Fractional reduction of peak current as a function of time spent at -140 mV, with superimposed double-exponential fits for the indicated concentrations of MgPpIX. (**F**, *top*) Time course of mean normalized peak currents at -30 mV with the application of 100 nM MgPpIX (blue). After about 100 s, the application pipette containing MgPpIX was withdrawn from the cell (light green bar), followed by washing with bath solution (green) and washing plus intermittent depolarizations (1-s episodes at -10 mV were inserted every 4 s between the test pulses to induce pronounced reverse use-dependence, dark green). (*bottom*) As in the top panel, but with the application of 10 µM of fatty-acid-free BSA, which accelerated the reversibility of MgPpIX-induced channel inhibition. All data in (B-F) are means ± sem, with *n* indicated in parentheses.

Both channel activation and inactivation contribute to the channel open probability in a voltage-dependent manner, making it difficult to determine which process underlies the reverse use-dependence. We therefore eliminated rapid channel inactivation by chemically modifying the inactivation motif. Such inactivation-deficient hNa_V_1.5 channels were still inhibited by MgPpIX, albeit to a lesser extent (Fig. EV2). The reverse use-dependence was directly observable as an increasing current during long depolarizations that opened the channels (Fig. EV2E). Although the coupling of channel activation and inactivation may be instrumental in the magnitude of the MgPpIX effect, channel inactivation is required neither for MgPpIX-induced channel inhibition nor for the reverse use-dependence.

### MgPpIX is a specific inhibitor of human Na_V_1.5 channels

To evaluate the potential specificity of MgPpIX for human Na_V_1.5, we also examined its effect on mouse Na_V_1.5 (mNa_V_1.5) as well as human Na_V_1.2, Na_V_1.4, and Na_V_1.7 expressed in HEK293T cells. MgPpIX (1 µM) noticeably inhibited mNa_V_1.5, while no such effect was detectable for the other isoforms (Fig. 3A,B). TTX-resistant hNa_V_1.8 channels, heterologously expressed in Neuro2A cells, were also unaffected by MgPpIX. The results underscore the remarkable specificity of MgPpIX for hNa_V_1.5 among the human isoforms. Assuming a Hill coefficient of unity, we estimate the IC_50_ values for hNa_V_1.2, hNa_V_1.4, and hNa_V_1.7 to be at least 100,000 times greater than that for hNa_V_1.5. Further, for direct comparison with hNa_V_1.5 with an IC_50_ ≈ 1 nM, we compiled a concentration–effect relationship for mNa_V_1.5 (Fig. EV3), yielding IC_50_ = 42.8 ± 5.0 nM and a Hill coefficient *n*_H_ = 0.81 ± 0.07. Thus, human Na_V_1.5 is about 40-fold more sensitive to MgPpIX than mouse Na_V_1.5.

**Figure 3.**
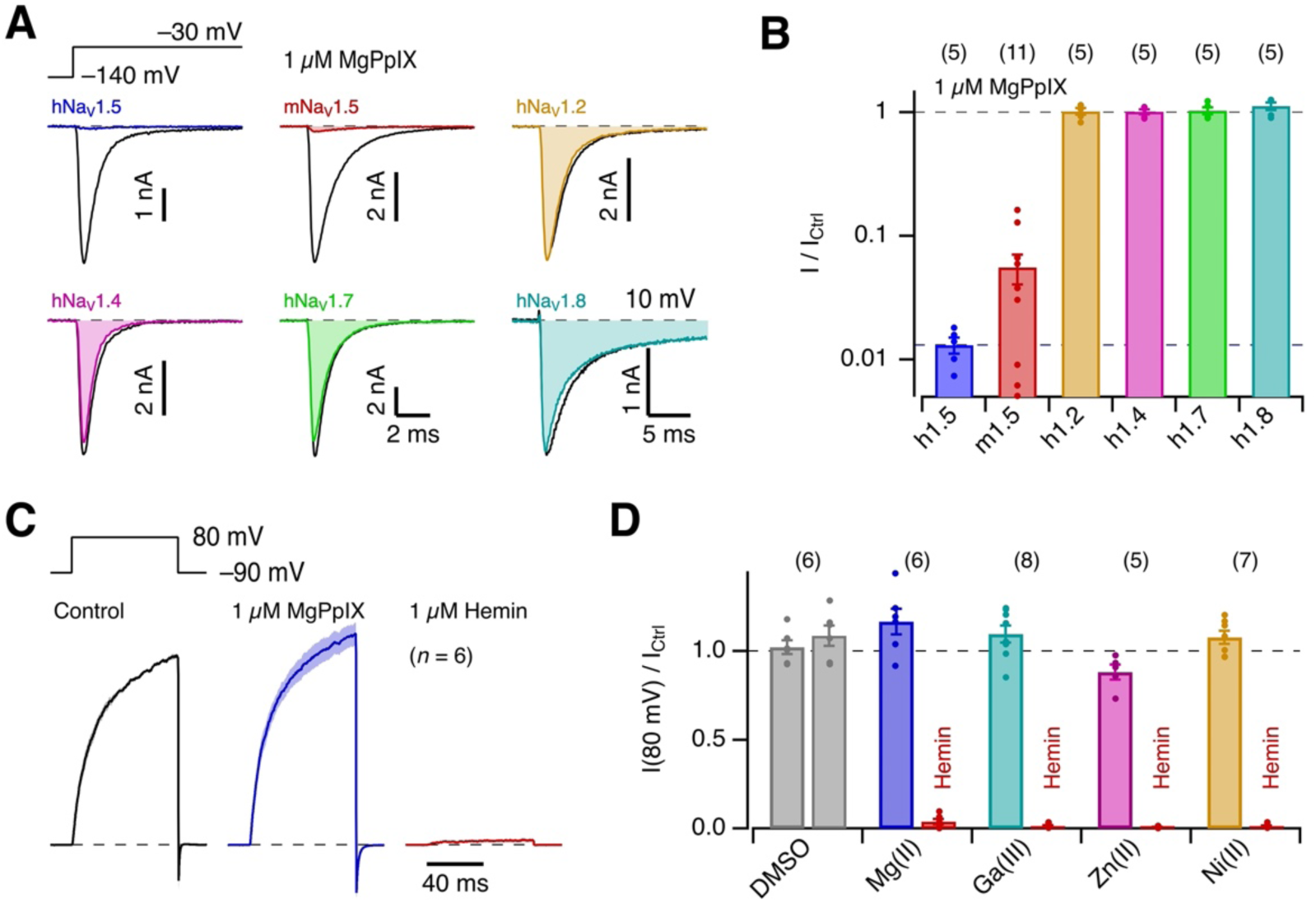
Specificity of MgPpIX for hNa_V_1.5 channels. (**A**) Representative whole-cell currents according to the indicated pulse protocol before (black) and about 5 min after the application of 1 µM MgPpIX (color) for hNa_V_1.5, mNa_V_1.5, hNa_V_1.2, hNa_V_1.4, hNa_V_1.7, and hNa_V_1.8. For hNa_V_1.8, depolarizing steps to 10 mV were applied. (**B**) Pooled results for the mean remaining relative current after 5 min of exposure to 1 µM MgPpIX for the indicated channel types. Data are means ± sem, with *n* in parentheses, and individual data points shown as dots. (**C**) Mean current traces with human BK_Ca_ currents in inside-out patches with 1 µM “intracellular” free Ca^2+^ in response to the indicated pulse protocol, normalized to the mean control current measured in the last 30 ms of the depolarization episode; sem is indicated in shading. Currents were measured before (black) and after the application of 1 µM MgPpIX (blue), as well as after a final application of 1 µM hemin (Fe(III)PpIX) to the intracellular side. (**D**) Current amplitudes at 80 mV relative to the control values for the application of the indicated MePpIX at 1 µM and the subsequent application of 1 µM hemin. Data are means ± sem with *n* in parentheses.

The impact of MgPpIX on hNa_V_1.5 channels appears to be unrelated to the previously reported influence of heme and hemin on ion channels with canonical heme-binding motifs. For example, the human large-conductance Ca^2+^-and depolarization-activated K^+^ channel (BK_Ca_), which is effectively inhibited by hemin when applied to the intracellular side of the plasma membrane (Tang *et al*., 2003), was not inhibited by 1 µM MgPpIX (Fig. 3C). Similarly, 1 µM GaPpIX and 1 µM NiPpIX did not block BK_Ca_ channels, while ZnPpIX had a minor effect (Fig. 3D) as previously reported (Tang *et al*., 2003).

### MgPpIX interacts with the voltage sensor of Na_V_-channel domain II

Despite hNa_V_1.5 being ∼40-fold more sensitive to MgPpIX, human and mouse Na_V_1.5 differ at only two residue positions in the extracellular loops in the voltage-sensor of domain II (VSD II): S743-A in the S1/S2 loop and S802-G in the S3/S4 loop (Fig. 4A,B). We found that the human-to-mouse mutation hNa_V_1.5-S802G in the S3/S4 loop diminished the sensitivity to 100 nM MgPpIX to a level identical to that of mNa_V_1.5 (Fig. 4C,D): the current remaining for hNa_V_1.5-S802G was 27.7 ± 4.1% (*n* = 5) as compared to 35.1 ± 2.8% (*n* = 9) for mNa_V_1.5 (t-test: *p* = 0.26). The mutation hNa_V_1.5-S743A did not affect the sensitivity of the channel to MgPpIX (remaining current 5.1 ± 1.1%, *n* = 7 as compared to 3.4 ± 0.4%, *n* = 10 for hNa_V_1.5, *p* = 0.30).

**Figure 4.**
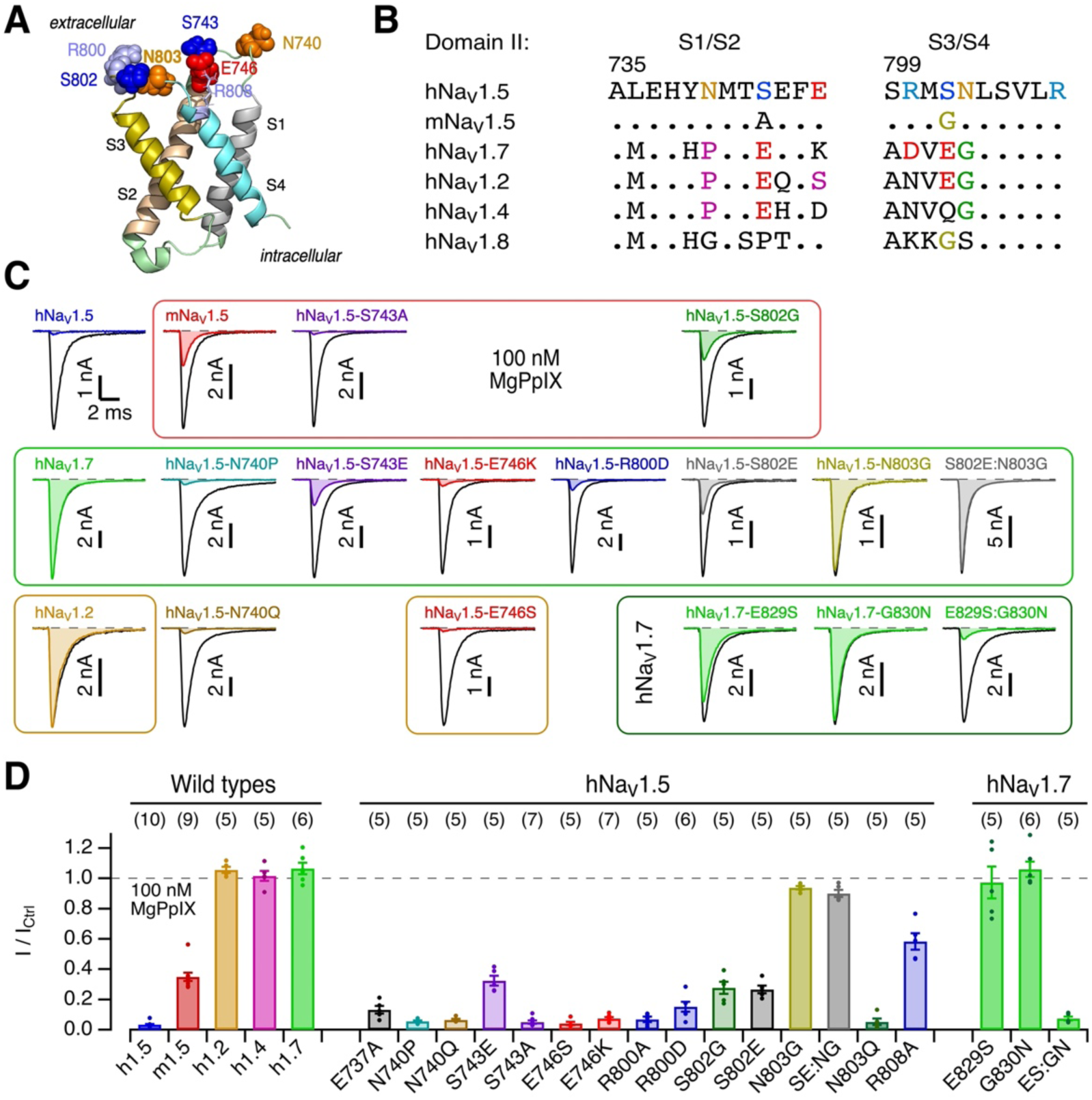
Mutagenesis of domain II voltage sensor. (**A**) Structure of the hNa_V_1.5 domain II voltage sensor (from PDB: 6LQA) with residues of interest highlighted. (**B**) Multiple sequence alignment of the domain II S1/S2 and S3/S4 loops. Residues identical to the hNa_V_1.5 sequence are presented as dots. (**C**) Representative current recordings at -30 mV for the indicated channel types and variants before (black) and 5 min after the application of 100 nM MgPpIX (colored). Red frame: hNa_V_1.5 mutations towards mNa_V_1.5; green frame: hNa_V_1.5 mutations towards hNa_V_1.7; brown frames: hNa_V_1.5 mutations towards hNa_V_1.2; dark green frame: hNa_V_1.7 mutations towards hNa_V_1.5. (**D**) Mean fractional peak current at -30 mV remaining 5 min after the application of 100 nM MgPpIX for the indicated channel types, as well as hNa_V_1.5- and hNa_V_1.7-based variants. Even at 10 nM, the inhibitory potency of MgPpIX was not markedly different between Na_V_1.5 (I/I_Ctrl_ 0.157 ± 0.028, *n* = 8) and hNa_V_1.7-E829S:G830N (0.16 ± 0.06, *n* = 4). For further functional properties of the mutants, see Suppl. Table 1 and Suppl. Fig. 3.

Among the human isoforms, hNa_V_1.5 is uniquely sensitive to MgPpIX. Yet, the extracellular loops in VSD II of hNa_V_1.5 differ from those of the MgPpIX-insensitive channels Na_V_1.2, Na_V_1.4, and Na_V_1.7 at only a few positions. Most prominently, these insensitive channels have in common a glycine residue at the equivalent position of hNa_V_1.5-N803 in the extracellular S3/S4 loop. In fact, the variant hNa_V_1.5-N803G was not inhibited by 100 nM MgPpIX (remaining current: 93.9 ± 1.2%, *n* = 5). The insensitivity arises from the presence of glycine at position 803 rather than a requirement for asparagine at that position for MgPpIX sensitivity; the variant hNa_V_1.5-N803Q was about as sensitive to MgPpIX as the wild type (Fig. 4D, Suppl. Fig. 2). Further mutational analysis (Suppl. Fig. 2) suggests that an amide group at position 803 is required for MgPpIX sensitivity. The mutation of serine at position 802 in hNa_V_1.5 to glutamate (hNa_V_1.5-S802E), as in Na_V_1.2 and Na_V_1.7, diminished the sensitivity slightly but not to the level of hNa_V_1.2 or hNa_V_1.7 (Fig. 4C,D). Thus, the high sensitivity of hNa_V_1.5 to MgPpIX compared with other hNa_V_ isoforms is attributable to the presence of N rather than G at position 803.

The importance of S802:N803 in the S3/S4 loop of VSD II for the sensitivity of hNa_V_1.5 to MgPpIX is corroborated by reverse mutagenesis of hNa_V_1.7, in which the equivalent residues, E829S and G830N in hNa_V_1.7, were changed to those in hNa_V_1.5. Substitution of either residue alone failed to confer high MgPpIX sensitivity. Concurrent substitutions (hNa_V_1.7-E829S:G830N) rendered hNa_V_1.7 highly sensitive: the remaining current in 100 nM MgPpIX was 7.5 ± 1.4% (*n* = 5). This sensitivity, while marked, is less than that of wild-type hNa_V_1.5, which is readily explained by other non-conserved residues in the S3/S4 loop (e.g., R800D) and the S1/S2 loop (e.g., N740P, S743E, and E746S); individual mutation of these residues in hNa_V_1.5 decreased its sensitivity to MgPpIX (Fig. 4D). Alteration of the charged residues E737 and R808, highly conserved among the hNa_V_ variants (Fig. 4B), also affected the channel’s response to MgPpIX (Fig. 4D).

### Structural modelling of MgPpIX interacting with hNa_V_1.5 in a lipid membrane

The mutagenesis results suggest that MgPpIX interacts with VSD II. Its impact on the channel is dramatically greater when the channel is deactivated or when the voltage sensors are probably in a “down” position. Structural assessment of a putative MgPpIX–channel interaction is challenging because the available cryo-electron microscopy structure of hNa_V_1.5 (PDB: 6LQA) shows all four voltage sensors in the activated “up” position (Fig. 5A, *left*). To circumvent this limitation, the hNa_V_1.5 VSD II was structurally aligned to the deactivated voltage sensor of K_V_7.1 (Mandala & MacKinnon, 2023). The resulting “down” structure as well as the original “up” position structure were embedded in membrane patches, each with one protein, and energy-minimized in short MD runs (Figs. 5A, EV4A,B).

**Figure 5.**
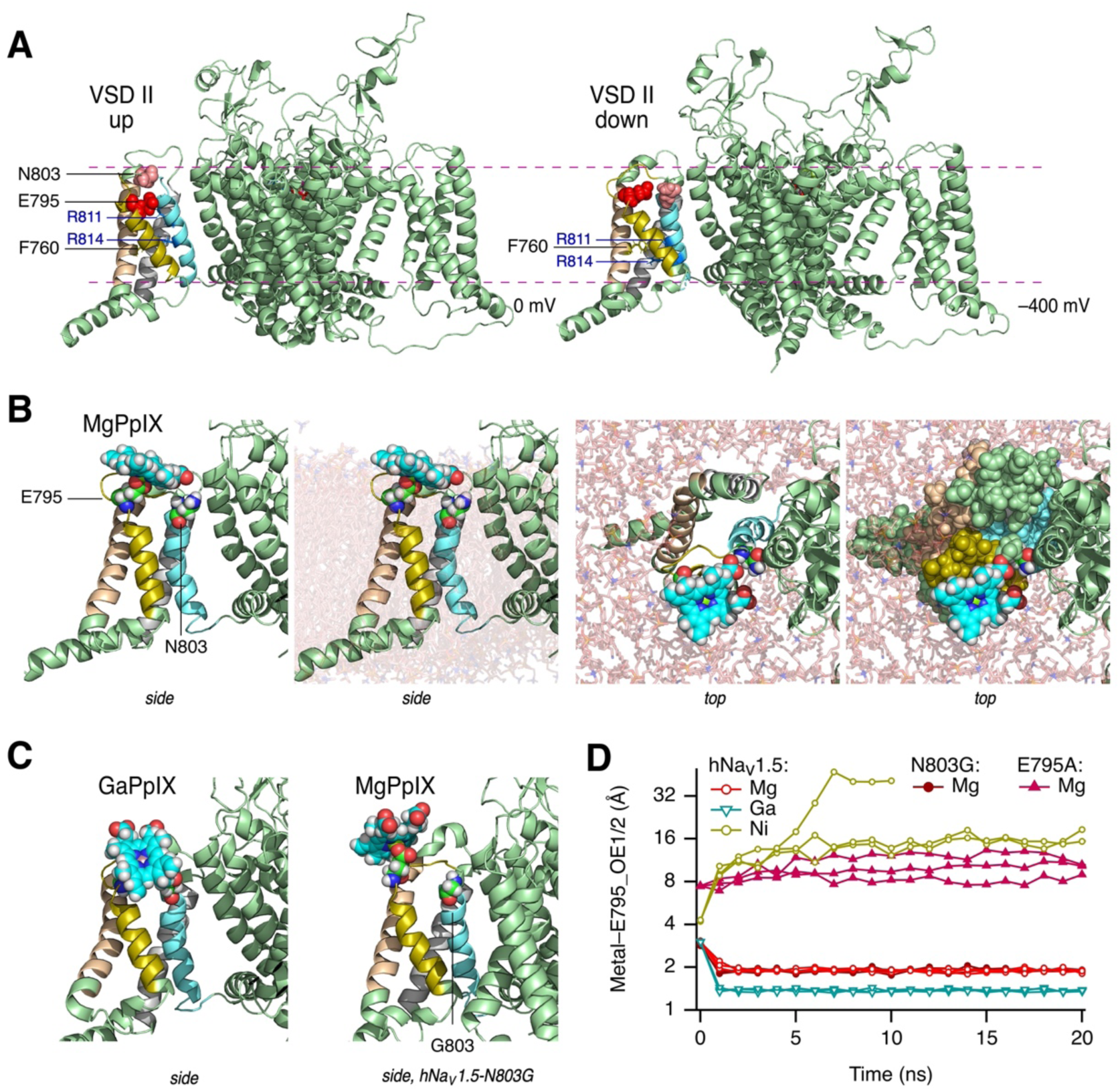
Structural model of hNa_V_1.5–MgPpIX interaction. (**A**) Structural model of hNa_V_1.5 based on a cryo-EM structure (PDB: 6LQA) with all voltage sensors being in its activated “up” position (*left*) or a structure in which the voltage sensor of domain II was brought in a deactivated or “down” position after alignment to K_V_7 channels (*right*). Residues E795 and N803 are shown as spheres, F760, R811, and R814 as sticks. S1-S4 are colored as in Fig. 4. Lipids, water, and ions are not shown for clarity. For the “up” configuration no transmembrane voltage was applied, for the “down” configuration simulations were performed with -400 mV. For details, see Fig. EV4. (**B**) Snapshots of 20-ns MD simulations of hNa_V_1.5 with VSD II in a “down” position, immersed in a lipid bilayer, and MgPpIX placed close to VSD II. The individual images show a zoom-up of VSD II of the same structure in different presentations: side view without lipids, side view with lipids, top view with lipids, and top view with lipids and all residues of VSD II as spheres. (**C**) As in panel (B, *left*) with GaPpIX docked to VSD II of the wild-type channel (*left*) and with MgPpIX docked to channel variant N803G (*right*). (**D**) Distances between the metal center of MgPpIX, GaPpIX, and NiPpIX to the minimum of the OE1 and OE2 oxygens of E975 in each of the three molecular dynamics runs in which the MePpIX was placed near E795 as a starting position. In addition, similar simulations with MgPpIX and the channel mutants E795A and N803G are included. For variant E795A, the distance of the Mg to the CA1 carbon of alanine is presented. For more examples and movies, see Figs. EV4, EV5, and Suppl. Movies.

In VSD II, residue F760 in S2 marks the charge transfer center (Tao *et al*, 2010), separating the electrically extracellular and intracellular compartments. In the “up” structure systems without any applied voltage (Fig. 5A, *left*), the S4 voltage-sensing charges R811 and R814 are extracellular to F760. R811 is located close to E737 in S1 and/or to E795 in the extracellularly facing S3/S4 loop (Fig. EV4B). In contrast, in the “down” structures simulated at an extreme negative voltage (Fig. 5a, *right*), R814 is intracellular to F760 and R811 is slightly intracellular to or at the level of F760 (Fig. EV4E), allowing the extracellular negative counter charge residues E737 and E795 to interact with water and ions in three of the four simulations (Fig. EV4). Salt bridges between E763 and R814 and possibly also between E737 and R800 are apparent in the “down” configuration (Fig. EV4F,H). These structural features are consistent with our current understanding of voltage-sensor dynamics.

Guided by the functional results that N803 in VSD II may be crucial for the MgPpIX–channel interaction, one MgPpIX molecule was manually placed near N803 on the extracellular side of the VSD II “down” structure. The resulting complex was energy minimized at the negative voltage. In one of the poses (Fig. 5B), MgPpIX was bound to the channel, stabilized by an electrostatic interaction between the magnesium and a side chain oxygen atom (OE1 or OE2) of E795, one of the S4 counter charge residues, with membrane lipids surrounding part of the protoporphyrin ring (Figs. 5B, EV5A). The propionate groups of MgPpIX sit just extracellular to N803. This binding pose was stable in all three independent simulations (Fig. 5D). The relative stability of different MePpIX was assessed by replacing the MgPpIX in this pose with GaPpIX or NiPpIX. GaPpIX remained with the channel like MgPpIX (Figs. 5C,D, EV5B). In contrast, NiPpIX dissociated away from the channel into the solvent or the membrane lipids in all three simulations (Fig. 5D, Suppl. Movies). The propensity of MgPpIX and GaPpIX to remain with the channel is consistent with their greater inhibitory potency on hNa_V_1.5 compared with NiPpIX (Fig. 1B). Further, the structural modeling results are in line with DFT-based calculations showing a more positive electrostatic potential at the metal center of MgPpIX and GaPpIX than that of NiPpIX (Figs. 1C, EV1).

The structural models in Fig. 5B-C suggest that the conserved residue E795 stabilizes the MgPpIX– channel interaction. In the hNa_V_1.5-E795A variant, 100 nM MgPpIX had no impact on the peak current at -30 mV (100.0 ± 2.3% remaining current, *n* = 5). Even 1 µM MgPpIX only partially inhibited the channels (57.2 ± 2.2% remaining current, *n* = 10) with no sign of reverse use-dependence (Suppl. Fig. 4). Furthermore, simulations using MgPpIX in hNa_V_1.5-E795A in the VSD II “down” conformation failed to show stable binding of MgPpIX (Fig. 5D, Suppl. Movies). Therefore, the functional and computational findings together validate our structural modeling approach.

The electrophysiological measurements showed that hNa_V_1.5-N803G is also resistant to 100 nM MgPpIX. In all three simulations of the N803G variant in the “down” conformation, MgPpIX remained bound via E795 (Figs. 5C,D, EV5C, Suppl. Movies). This finding suggests that the mutation N803G may interfere with the association step and/or efficacy.

### Impact of MgPpIX on *SCN5A*-expressing human cancer cell lines

In multiple human breast cancer cell lines, endogenous expression of *SCN5A* is linked to malignancy, likely due to functional hNa_V_1.5 channels in the plasma membrane. Interference with the channel function has been suggested to diminish cell migration and invasiveness (Brackenbury *et al*, 2007; Driffort *et al*, 2014; Lee *et al*, 2019; Leslie *et al*, 2024; Onkal & Djamgoz, 2009). We thus examined the impact of various MePpIX on cell migration of the breast cancer cell line MDA-MB-231 using a wound-healing assay. To minimize confounding with cell proliferation, the medium contained only 2% FBS. Under the control condition, the scratch-induced gap narrowed within 4 h due to cell migration. With MgPpIX (100 nM), the gap narrowing was reduced, suggesting delayed cell migration (Fig. 6A,B). The vehicle (DMSO), as well as PpIX, NiPpIX, or SnPpIX, which did not affect hNa_V_1.5 function (Fig. 1), failed to influence gap closure: the gap became narrower as in the control condition. FePpIX, GaPpIX, and ZnPpIX diminished gap closure, albeit less potently than MgPpIX (Fig. 6B). These results, which show a correlation between the MePpIXs’ potency on hNa_V_1.5 and cell migration, were corroborated by a transwell migration assay with overnight exposure to MgPpIX. Using a chemoattractant gradient from 0% (insert) to 2% FBS (wells), the number of migrated cells with 100 nM MgPpIX in the insert was reduced compared to control, PpIX, and SnPpIX application (Fig. 6C,D).

**Figure 6.**
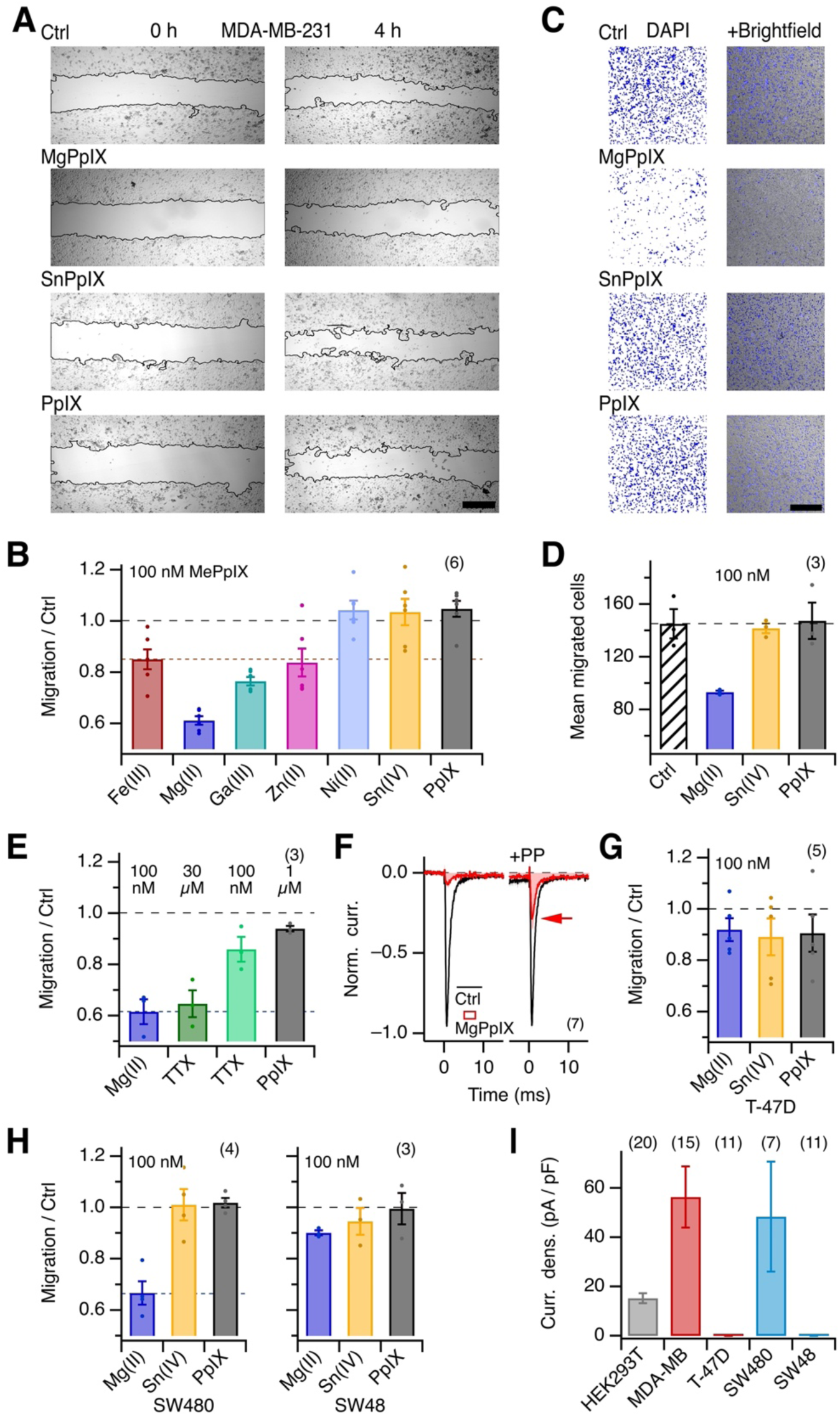
MgPpIX slows down the migration of cancer cells expressing *SCN5A*. (**A**) Representative images of MDA-MB-231 cell cultures right after introducing a scratch gap (0 h) and 4 h later. After scratch introduction, the cells were kept in control medium with DMSO (Ctrl) or 100 nM of the indicated MePpIX, dissolved in DMSO. The cells were kept in the dark to prevent phototoxicity induced by the MePpIXs. Continuous black lines indicate the automatically detected scratch borders. Scale bar: 400 µm. (**B**) Mean gap-closure migration of MDA-MB-231 cells (relative to the migration under control conditions, i.e., in the presence of the vehicle DMSO) in the presence of 100 nM of PpIX or the indicated MePpIX variants. (**C**) Representative confocal images of DAPI-stained MDA-MB-231 cell nuclei detected on the trans side in a transwell migration assay with an 8 µm pore size and a chemoattraction gradient from 0% (insert) to 2% FBS (wells); background in white. The panels on the right show superpositions of the DAPI staining (blue) with transmission images. Cells were kept overnight under control conditions or in the presence of 100 nM of the indicated MePpIX. Images were taken through a 10x objective with a 3x zoom factor. Cell counting was performed based on the DAPI staining using ImageJ software. Scale bar: 400 µm. (**D**) Mean number of migrated cells from experiments as shown in (C). (**E**) Mean migration in a wound-healing assay of MDA-MB-231 cells over 4 h in the presence of the indicated concentrations of MgPpIX, TTX, and PpIX. (**F**) Mean normalized current traces obtained from MDA-MB-231 cells under voltage-clamp conditions with step depolarizations to -30 mV without (black) and with the application of 100 nM MgPpIX. The currents on the right followed an episode of 250 ms at 40 mV to highlight the reverse use-dependence of these Na_V_ channels. (**G**) Mean relative migration in a wound-healing assay (as in (B) and (E)) but for T-47D cells. (**H**) As in (G) but for the colon carcinoma cell lines SW480 and SW48. (**I**) Mean current density, based on the peak currents at -30 mV, for HEK293T cells and the indicated breast cancer (MDA-MB-231, T-47D) and colon carcinoma cells (SW480, SW48). Data bars are means ± sem with *n* in parentheses; in panels B, D, E, G, and H, the number of independent cell batches is given.

That Na_V_ channel inhibition underlies the MgPpIX-induced reduction in cell migration is supported by wound-healing assays comparing the effects of TTX. At 30 µM, a concentration high enough to block even hNa_V_1.5 channels, TTX had a similar inhibitory impact on gap closure as 100 nM MgPpIX, while at 100 nM, TTX had only a weak effect (Fig. 6E). The results thus collectively show that MgPpIX inhibits cell migration with much greater apparent efficacy than TTX and that MDA-MB-231 cells primarily harbor MgPpIX-sensitive but only weakly TTX-sensitive Na_V_ channels. This idea is further supported by direct electrophysiological measurements of endogenous Na_V_-channel current in MDA-MB-231 cells (mean current density at -30 mV: 56.3 ± 12.4 pA/pF, *n* = 15). MgPpIX (100 nM) decreased the current to 6.8 ± 0.7% and 100 nM TTX slightly diminished the current further to 3.5 ± 0.5%. The neonatal splice variant of hNa_V_1.5 (nhNa_V_1.5), which is expressed in MDA-MB-231 cells (Brackenbury *et al*., 2007), was as sensitive to MgPpIX as the adult variant (Suppl. Table 1).

While the pronounced MgPpIX sensitivity in MDA-MB-231 cells highlights the importance of hNa_V_1.5, not all breast cancer lines possess hNa_V_1.5 channels. For example, we found no Na_V_ current in the human breast cancer cell line T-47D (*n* = 11), and its cell migration was unaffected by MgPpIX (Fig. 6G). A similar correlation between the impact of MgPpIX on cell migration and the presence of functional hNa_V_1.5 channels was observed for two human colorectal carcinoma cell lines: SW480 and SW48. SW480 cells showed an Na_V_ current density of 48.4 ± 22.3 pA/pF, which was decreased to only 13.5 ± 4.1% (*n* = 7) by 100 nM MgPpIX; their cell migration was also inhibited (Fig. 6H,I). In contrast, SW48 cells have no hNa_V_1.5 channels (*n* = 11), and their migration was unaffected by MgPpIX (Fig. 6H,I). Furthermore, neither SnPpIX nor PpIX had any impact on the migration of SW480 or SW48 cells (Fig. 6H,I).

## Discussion

The study here shows that Mg-protoporphyrin IX, a metal protoporphyrin intermediate in the plant chlorophyll synthesis pathway between protoporphyrin IX and MgPpIX monomethylester, is by far the most potent and specific inhibitor of human Na_V_1.5 channels with an IC_50_ of about 1 nM. Notably, MgPpIX occurs in plants at concentrations of 30 nM (Mochizuki *et al*., 2008). The efficacy of inhibiting hNa_V_1.5 currents depends on the metal atom; Mg(II)PpIX is about 100-fold more effective than Fe(III)PpIX, whereas Ni(II)PpIX, Cu(II)PpIX, and Sn(IV)PpIX are completely ineffective. Quantum mechanical calculations with MePpIX variants reveal that an electropositive surface potential of the metal is necessary for potent inhibition (Fig. EV1), and MgPpIX has the most electropositive metal center among the MePpIX variants examined. While Mg(II)PpIX, Ni(II)PpIX, and Cu(II)PpIX contain a metal center that typically forms divalent ions in aqueous solution, the electropositive surface potential in the PpIX scaffold is markedly greater for magnesium. Even Fe(III)PpIX, with a metal center typically forming a trivalent cation, shows a weaker electropositive surface potential than Mg(II)PpIX. The physicochemical interaction of MgPpIX with hNa_V_1.5 channel proteins, where the metal center electropositive surface potential plays a critical role, represents a fundamentally distinct paradigm of interaction compared with the previously reported heme or hemin interactions with other ion channels (Burton *et al*, 2016; Sahoo *et al*, 2013; Sahoo *et al*, 2022; Tang *et al*., 2003).

In addition to the aforementioned striking role of the metal center, another unique feature of the interaction of MgPpIX with hNa_V_1.5 is the pronounced reverse use-dependence; channel inhibition is alleviated by channel activation. This characteristic identifies MgPpIX as a gating modifier of hNa_V_1.5 channels, functionally resembling select natural peptide toxins. The µO-conotoxin MrVIA from marine cone snails inhibits Na_V_ channels in reverse use-dependent manner (Leipold *et al*, 2007), and the impact of the scorpion µ toxin Tz1 on Na_V_1.4 is markedly diminished by activating prepulses (Leipold *et al*, 2012). In both cases, the dependence is conferred by VSD II. This is also the case for MgPpIX: we identified residues E795 and N803 in VSD II of hNa_V_1.5 as critical determinants for MgPpIX activity on cardiac Na_V_ channels. The VSD II-dependent inhibition of action of hNa_V_1.5 by MgPpIX is reminiscent of the action of suzetrigine, a small-molecule analgesic that inhibits neuronal hNa_V_1.8 targeting its VSD II (Jo *et al*, 2025; Osteen *et al*., 2025). Collectively, VSD II may represent a promising structural target for isoform-specific modulation of Na_V_ channels.

The mutagenesis results suggest that MgPpIX interacts with the extracellularly accessible loops of VSD II. One critical residue is N803 in hNa_V_1.5, which corresponds to a glycine at the equivalent position in other MgPpIX-insensitive Na_V_ isoforms. In addition, neighboring residues in the S1/S2 and S3/S4 loops contribute to the full activity as indicated by the difference between mouse and human Na_V_1.5 (hNa_V_1.5-S802G) and the reverse mutagenesis of hNa_V_1.7 toward hNa_V_1.5 (hNa_V_1.7-E829S:G830N), which conferred MgPpIX sensitivity.

Asparagine at position 803 is not the only amino acid capable of supporting the full sensitivity to MgPpIX. Similar high sensitivity is observed in the hNa_V_1.5 variants with Q, H, R, and K at that position (Suppl. Fig. 2). Thus, an amide group at position 803 may be required for full MgPpIX sensitivity. The apparent importance of an amide side chain is not unprecedented. Although structurally quite dissimilar, a binding of MgPpIX to residue N211 in Gun4-1, a plant porphyrin-binding protein, has been reported (Kopečná *et al*, 2015). In addition to protein interactions, the amphipathic character of MgPpIX suggests that the lipids surrounding the channel protein may contribute to its inhibitory action. Recovery from inhibition is very slow and incomplete; near full recovery could only be achieved by washing with fatty-acid-free BSA, indicative of a strong interaction of MgPpIX with the lipids and/or with highly hydrophobic parts of the channel.

The strong reverse use-dependence of the MgPpIX action indicates that the binding pose of MgPpIX is critically dependent on the voltage-dependent conformational changes of hNa_V_1.5. Since an hNa_V_1.5 structure featuring the voltage sensors in a deactivated “down” configuration is currently unavailable, we created a model, based on the hNa_V_1.5 cryo-EM structure, in which VSD II was aligned with the “down” position of K_V_7.1 channels. This structure, incorporated into a lipid membrane with a negative membrane potential, was subjected to MD simulations with MgPpIX and other MePpIX placed in the vicinity of VSD II (Figs. 5, EV5). While not definitive or exclusive, the structural modeling results suggest the following mechanism to account for the use-dependent inhibition. When VSD II is deactivated or “down” at negative voltages, the VSD-II voltage-sensing charges such as R811 and R814 are located intracellular to the VSD-II charge transfer center F760, leaving the extracellular-facing negative counter charges such as E795 to interact with ions, water, and MgPpIX. The negative side chain oxygens of E795 electrostatically interact with the electropositive metal center of MgPpIX while the hydrophobic protoporphyrin macrocycle interacts with the surrounding membrane lipids. In this manner, MgPpIX stabilizes the deactivated “down” conformation of VSD II and the channel remains closed and inhibited. This interaction is disrupted when VSD II is activated by strong depolarization, driving R814 and R811 outward to interact with extracellular negative counter charges such as E795. As a result, stable binding of MgPpIX is no longer possible and the channel inhibition is relieved. The variant hNa_V_1.5-N803G (Figs. 5C,D, EV5) is functionally insensitive to MgPpIX; however, the structural modeling here shows a stable interaction with MgPpIX. This discordance probably suggests that while MgPpIX may engage with VSD II even in the absence of asparagine at position 803, only with N803 does the interaction impede the movement of VSD II from the resting “down” to the activated “up” state. Altogether, the use-dependent inhibition arises from the voltage-dependent movement of VSD II and its influence on the accessibility of the negative counter charges such as E795 for the metal center of MgPpIX. While our structural modeling has identified a plausible pose of MgPpIX binding involving E795 in VSD II, it is important to note that the results here do not rule out the existence of other interaction arrangements.

Functional expression of Na_V_1.5 channels in breast, colon, and several other cancer cell types is associated with a more invasive metastatic phenotype, and channel inhibition *in vitro* can suppress this invasive activity. The unmatched potency and subtype selectivity of MgPpIX as inhibitor of Na_V_1.5 currents renders it a promising lead structure for the development of new pharmacological tools to improve cancer treatment. However, the direct translation of MgPpIX as an anti-tumor agent may be considered challenging because of the critical role of Na_V_1.5 channels in cardiac myocytes, where a high risk of cardiac arrhythmias looms. Nevertheless, a possible therapeutic window in cancer cells likely exists because of the reverse use-dependence of MgPpIX. Strong depolarization, as during rhythmic cardiac action potentials, should diminish the inhibitory potency of MgPpIX, thus diminishing the cardiac arrhythmia risk. In contrast, potent inhibition should prevail at typical negative resting potentials of cancer cells. It is noteworthy that MgPpIX is a natural metabolite found in plants. Further, the potential of using a Na_V_1.5 inhibitor as an anti-tumor agent has been suggested by Driffort *et al*. (Driffort *et al*., 2014), who demonstrated that in a mouse model with xenografted Na_V_1.5-expressing human breast cancer cells, the Na_V_1.5 blocker ranolazine was well-tolerated and led to reduced lung colonization by the cancer cells.

In summary, MgPpIX is a natural compound with unmatched potency and specificity as an inhibitory modulator of hNa_V_1.5 channels. The inhibitory action is mediated through the voltage sensor in domain II, which is trapped by MgPpIX in a resting configuration. This unique mechanism of action, along with the role of hNa_V_1.5 in multiple cancers, positions MgPpIX as a promising lead compound for the development of anti-tumor therapeutic strategies.

## Methods

### Electrostatic surface potentials

Structures and electrostatic potential maps of various MePpIX were calculated using GaussView 6.0.16/Gaussian 16 (Gaussian, Wallingford, CT, USA), running on a Linux workstation with 20 CPU cores. MePpIX structures were optimized using DFT (density functional theory) calculations at the B3LYP level, employing the aug-cc-pVDZ basis set and water as a solvent under a self-consistent reaction field (SCRF) and default or loose convergence criteria, as specified in Fig. EV1. A low-spin state was assumed for all structures. For each MePpIX, we generated one structure in which the carboxylate groups were neutralized by placing one H^+^ ion between the carboxylate oxygens. The surplus of positive charge of the trivalent metal ions was compensated by placing one Cl^−^ ion close to the metal.

The final electrostatic surface potential (ESP) maps were generated in GaussView by mapping onto electron density isosurfaces of 0.0004 atomic units, and the energies were converted from Hartree to electron volts (eV: 1 Hartree = 27.2114 eV) for display purposes. ESP-derived atomic charges were estimated for the optimized structures using the Merz-Singh-Kollman (MK) population analysis and UFF (Universal Force Field) radii (Iop 6/41=10, 6/42=10). ESP points were sampled from 10 layers with a density of 10 points per unit area, and the layers were defined based on UFF radii.

### Protein structure modeling

The cryogenic electron microscopy (cryo-EM) structure of hNa_V_1.5 (PDB: 6LQA) shows the voltage-sensor domains in their activated “up” conformation (Li *et al*., 2021). To infer the structure of hNa_V_1.5 with voltage-sensor domain II in the resting “down” conformation, S1-S4 of domain II (residues 717-819) in hNa_V_1.5 was modelled using the cryo-EM structure of K_V_7.1 obtained in hyperpolarized lipid vesicles (KCNQ1; PDB: 8SIN) according to I-TASSER (Yang & Zhang, 2015). The resulting homology model structure was energy optimized using short all-atom molecular dynamics simulation runs using CHARMM-GUI (Wu *et al*, 2014) and CUDA-enabled NAMD2/3 with the CHARMM36m force field. Briefly, the channel complex was placed in a rectangular membrane patch composed of cholesterol (20%), dipalmitoylphosphatidylcholine (DPPC, 40%) and dioleoylphosphatidylcholine (DOPC, 40%) in the extracellular leaflet and cholesterol (20%), DPPC (25%), DOPC (25%), dipalmitoylphosphatidylethanolamine (DPPE, 10%), dioleoylphosphatidylethanolamine (DOPE, 10%), dipalmitoylphosphatidylserine (DPPS, 5%), and dioleoylphosphatidylserine (DOPS, 5%) in the intracellular leaflet (van Meer *et al*, 2008) in the presence of 150 mM KCl. The periodic boundary simulation box was about 180 Å x 180 Å x 130 Å in size (Fig. EV4A). The simulation systems were equilibrated according to the default parameter values suggested by CHARMM-GUI and the simulations were performed in the NPT ensemble at 30°C and 1 atm. For each condition studied, three independent systems were typically generated and simulated for at least 20 ns. The simulations of hNa_V_1.5 with the activated voltage sensors (PDB: 6LQA) were performed without applied voltage, while for the simulations of the models with voltage-sensor domain II in the resting “down” conformation, a saturating membrane potential of about - 400 mV was applied using the NAMD eField command. In the structure of hNa_V_1.5 (PDB: 6LQA), some loop segments are unresolved; thus, in the simulations, hNa_V_1.5 was modeled as a trimeric complex. The mutations N803G and E795A were made in CHARMM-GUI.

To simulate the interactions of hNav1.5 with MgPpIX, GaPpIX or NiPpIX, the following approach was used. The atomic coordinates and the partial charges were obtained from the Gaussian-optimized files (see above) and were used to generate the topology (.rtf) and the parameter (.prm) files using those for heme available from CHARMM-GUI as the templates. Because the bond parameters for these MePpIX compounds were not experimentally verified in this study, the results should be taken as approximations. Guided by the functional results that N803 in VSD II may be crucial for the MgPpIX-channel interaction, one MgPpIX was placed near N803 on the extracellular side of the voltage sensor as judged by eye. Six of such poses were energy minimized using short molecular dynamics simulations as described. By the end of the simulation, in all but one system, MgPpIX moved away from the channel to the extracellular solution or to the membrane lipids. In one system, MgPpIX remained near the channel and the results from this pose are presented (Figs. 5B,D, EV5). To compare the relative stability of NiPpIX and GaPpIX, the MgPpIX in this stably bound pose was replaced with a NiPpIX or GaPpIX molecule and the modified systems were energy minimized. For each of MgPpIX, NiPpIX and GaPpIX, three independent systems were studied and produced similar results. A similar binding-pose of MgPpIX was examined for variants N803G and E795A.

Images of protein structure were rendered with the PyMOL Molecular Graphics System (Version 3.0 Schrödinger, LLC.)

### Channel-coding DNA plasmids

The coding sequence of human Na_V_1.5 (hNa_V_1.5, *SCN5A*, Acc. no. M77235) was inserted into pRC/CMV vector plasmids and expressed in HEK293T cells as described previously (Chen & Heinemann, 2001). The following Na_V_ constructs were used in a similar manner: mNa_V_1.5 (mouse, AJ271477.1), hNa_V_1.2 (NM001040143), hNa_V_1.4 (AAO83647), hNa_V_1.7 (NP002968), and hNa_V_1.8 (Q9Y5Y9.2). The neonatal splice variant of hNa_V_1.5 (nhNa_V_1.5) was used as described by Schroeter *et al*. (Schroeter *et al*, 2010). Recombinant DNAs for all Na_V_ subgroups were engineered using PCR-based methods and verified by DNA sequencing. For the functional evaluation of BK_Ca_ channels (K_Ca_1.1), we used human *KCNMA1* (Acc. no. U11058).

### Cell culture and transfection

Human embryonic kidney 293T cells (HEK293T, from CAMR; Porton Down, Salisbury, UK) were cultured in Dulbecco’s Modified Eagle’s Medium (DMEM) with Nutrient Mixture F-12 (Gibco). Mouse neuroblastoma cells (Neuro-2A, DSMZ, Braunschweig, Germany) were cultured in DMEM. The breast cancer cell lines MDA-MB-231 and T-47D were cultured in RPMI 1640 medium (Sigma, R7388), while the colorectal cancer cell lines SW480 and SW48 were maintained in DMEM. Media were supplemented with 10% FBS, and the cells were incubated at 37°C in a humidified atmosphere containing 5% or 10% (Neuro-2A) CO_2_.

Cells were transfected with the channel-coding plasmids using ROTI®Fect (Carl Roth, Karlsruhe, Germany). Visualization of transfected cells was achieved by co-expressing green fluorescent protein or in the case of BK_Ca_ channels, CD8+ with subsequent use of anti-CD8 coated microbeads. Neuro-2A cells were utilized for the expression of hNa_V_1.8 channels; all other constructs were expressed in HEK293T cells. Ion currents were recorded 1-2 days after transfection.

### Wound-healing assay

Cells (50,000 to 70,000 per well) were seeded in a 48-well plate (Greiner Bio-One, Frickenhausen, Germany) and cultured for one day in their respective culture media supplemented with 10% FBS. Upon achieving approximately 90% confluency, a 200 µl sterile pipette tip was used to create a linear scratch (wound) in each well. Subsequently, the cells were rinsed with phosphate-buffered saline (PBS) and the medium was replaced with the same formulation containing only 2% FBS to minimize cell proliferation and to emphasize cell migration. Following the medium change, compounds were added to the wells and incubated for 4 h after treatment. To prevent light-induced degradation of the MePpIX variants and a photodynamic effect on the cells, the plates were protected from light. Images were captured before treatment and after 4 h of treatment using a live-cell imaging inverted microscope (Nikon Eclipse-Ti, USA) equipped with a DS-Qi2 camera (14-bit, Nikon) and an X-Cite 120 LED light source (Excelitas Technologies, USA), all managed using NIS-Elements software version 4.6 and temperature-controlled by an Okolab unit (Okolab s.r.l., Pozzuoli, Italy). Images were analyzed with ImageJ (NIH, Bethesda, MD, USA) using the “wound_healing_size_tool_updated” plugin (Suarez-Arnedo *et al*, 2020) to quantify wound areas.

### Transwell migration assay

The transwell migration assay was conducted to evaluate cell migration through the pores, facilitated by a chemotactic gradient. Cell suspension (100 µl) containing 50,000 cells in FBS-free medium was placed in the upper side of an insert with 8 µm pores (CellQART, Northeim, Germany), situated in 750 µl of medium supplemented with 2% FBS in a 24-well plate to establish a chemotactic gradient. For the treatment phase, 100 µl of various compounds in FBS-free medium was introduced into each well, mixed with the suspended cells, and the assembly was covered with aluminum foil to protect it from light. 24 h later, the inserts containing cells were fixed using 3.7% formaldehyde in PBS, permeabilized with methanol, and stained with DAPI (AppliChem, Darmstadt, Germany). Non-migrated cells on the inner surface of the insert were mechanically removed. Cells that had migrated to the outer side of the chamber were visualized, and images were captured using a LEICA DMi8 TCS SP8 inverted laser-scanning microscope operated with the Leica Application Suite X (Leica Biosystems, Wetzlar, Germany). Migrated cells were counted with ImageJ.

### Electrophysiological recording

Na_V_ currents were measured in the whole-cell configuration of the patch-clamp method with an EPC10 amplifier and PatchMaster acquisition software (HEKA Elektronik, Lambrecht, Germany). Pipettes were fabricated from borosilicate glass with a filament; the tips were coated with dental wax and fire-polished to yield resistances between 0.8 and 1.5 MΩ. Current was sampled at a rate of 50 kHz with a 10-kHz low-pass antialiasing filter. Only recording configurations with a series resistance below 3 MΩ were used; the series resistance was corrected electronically up to 75%. Leak and capacitive currents were subtracted using a p/6 correction protocol. The junction potential of about 10 mV was not corrected. The holding membrane voltage was -120 mV; before test pulses the cell was hyperpolarized to -140 mV for 50 ms to ensure full recovery from fast inactivation. Data recording only started after more than 8 min of equilibration of the cell at -120 mV in the whole-cell configuration. Prior to analysis and display, current traces were Gaussian-filtered with a cutoff frequency of 4 kHz.

For voltage-clamp recordings, the pipette solution was (in mM): 105 CsF, 35 NaCl, 10 ethylene glycol tetraacetic acid (EGTA), and 10 4-(2-hydroxyethyl)-1-piperazineethanesulfonic acid (HEPES), with the pH adjusted to 7.4 (CsOH). The bath solution contained 146 NaCl, 2 CaCl_2_, 2 MgCl_2_, 4 KCl, and 10 HEPES, with a pH of 7.4 (NaOH). For recordings of hNa_V_1.8 currents in Neuro-2A cells, the bath solution also included 1 µM TTX to block Na_V_ currents that are endogenous to Neuro-2A.

Currents of BK_Ca_ channels were measured in inside-out patches after expression in HEK293T cells, using an EPC10 amplifier (HEKA Elektronik) and PACHMASTER NEXT acquisition software (Multi-Channel Systems GmbH, MCS, Reutlingen, Germany). The holding voltage was -90 mV, and currents were recorded for depolarization steps up to 80 mV. Pipettes with resistances of 1.8-2.5 MΩ were used. The extracellular (pipette) solution contained (in mM): 138 NaCl, 4 KCl, 2 CaCl_2_, 2 MgCl_2_, and 10 HEPES, pH 7.4 (NaOH); the intracellular (bath) solution with about 1 µM free Ca^2+^ contained: 135 KCl, 8.63 CaCl_2_, 2 MgCl_2_, 10 EGTA, and 10 HEPES, pH 7.4 (KOH).

All recordings were obtained at room temperature (20-23°C). Metal protoporphyrins, diluted from stock solutions in DMSO, were applied by adding 1/3 of the final bath volume and mixing. Transmission and fluorescence light sources were turned off during data acquisition.

*Current-voltage relationships.* Currents were elicited with depolarizing steps from -100 to 60 mV in increments of 10 mV. The respective peak currents *I*(*V*) were fitted assuming a linear single-channel characteristic and *m*=3 independent activation gates according to Hodgkin and Huxley:

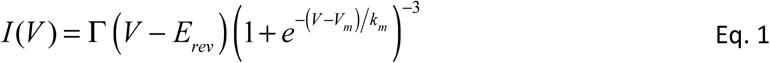

with the total conductance Γ, the reversal potential *E*_rev_, the half-maximal voltage of gate activation *V*_m_, and the slope factor *k*_m_, the latter characterizing the voltage dependence.

*Voltage dependence of fast inactivation.* The voltage dependence of fast inactivation was measured using a two-pulse protocol with depolarizations to -10 mV, separated by 500-ms intervals at variable voltage, *V*. The normalized peak currents were fitted with a Boltzmann-type function,

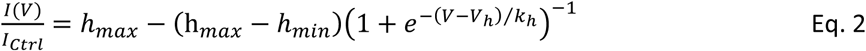

yielding the half-maximal voltage of inactivation *V*_h_, and the slope factor *k*_h_ characterizing the voltage dependence; *h*_min_ and *h*_max_ denote the minimal and maximal channel availability, respectively.

All variants of hNa_V_1.5 and hNa_V_1.7 produced voltage-gated Na_V_ currents of approximately equal amplitude and voltage dependencies of activation and inactivation, as shown in Suppl. Fig. 3 and Suppl. Table 1.

*Assessment of the reverse use-dependence.* The fractional amount of current relief from inhibition as a function of the voltage of a 500-ms prepulse (+PP) (Fig. 2C) was described by the following Boltzmann-type function:

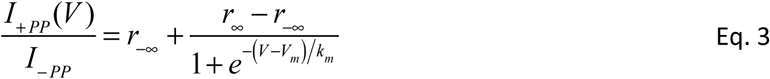

with the peak current before the prepulse (*I*_–PP_), the peak current after the prepulse (*I*_+PP_), the minimum and maximum ratio (*r*_οο_, *r*_–οο_), the half-maximal voltage of activation *V*_m_, and the slope factor *k*_m_.

The kinetics of block relief as a function of time at the depolarizing voltage of 50 mV (Fig. 2D) and of block establishment at –140 mV (Figs. 2E, EV2F, EV3E) was described by double-exponential functions.

### Chemicals

Protoporphyrin IX (PpIX, P8293) and hemin (Fe(III)PpIX, 51280) were acquired from Sigma-Aldrich. Additional metal-protoporphyrins, including magnesium-PpIX (Mg(II)PpIX, P10270), gallium-PpIX (Ga(III)PpIX, P40467), zinc-PpIX (Zn(II)PpIX, Zn625-9), manganese-PpIX (Mn(III)PpIX, P562-9), nickel-PpIX (Ni(II)PpIX, P40192), copper-PpIX (Cu(II)PpIX, P40769), and tin-PpIX (Sn(IV)PpIX, Sn749-9), were obtained from Frontiers Specialty Chemicals (Logan, UT, USA). Stock solutions of these compounds were prepared at 1 mM in dimethyl sulfoxide (DMSO, Sigma). Aliquots of these solutions were stored in at -20°C to preserve their stability for future experiments. MePpIX were dissolved in the bath solution, briefly vortexed, and applied to the recording chamber with gentle mixing.

### Data analysis

Data were analyzed using FITMASTER NEXT (MCS) and Igor Pro 9 (WaveMetrics, Lake Oswego, OR, USA) software. Averaged data are presented as means ± sem, individual data points are shown for *n* ≤ 10. The following statistical tests were used: t-test for normally distributed data and Wilcoxon rank tests otherwise. The resulting two-sided *p* values are provided as data descriptors.

The concentration dependence of compound-induced current inhibition was described as follows,

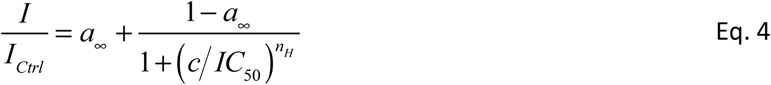

with the compound concentration *c*, the half-inhibitory constant *IC*_50_, the Hill coefficient *n*_H_, and a residual non-blocked fraction *a*_οο_.

## Abbreviations

BK_Ca_: large-conductance Ca^2+^-and depolarization-activated K^+^ channel
DFT: density functional theory
DTT: dithiothreitol
DMEM: Dulbeccós modified media
DMSO: dimethyl sulfoxide
ESP: electrostatic surface potential
FBS: fetal bovine serum
IC_50_: half-inhibitory concentration
MD: molecular dynamics
MePpIX: metal protoporphyrin
Na_V_: voltage-gated sodium channel
PpIX: protoporphyrin IX
TTX: tetrodotoxin
VSD: voltage-sensor domain

## Acknowledgments

Professors Th. Zimmer, Jena, and E. Leipold, Lübeck, for providing the channel-coding DNA plasmids.

## Research Funding

Support by the German Research Foundation (DFG, SHH: HE2993/18-1) and the National Institutes of Health (NIH, TH: GM121375).

## Author Contributions

MJ, mutant generation, electrophysiological recordings, cell-biological assays, data analysis; MA, electrophysiological recordings and quantum-chemical calculations; AB, JR, and GG, electrophysiological recordings; RS, molecular biology of expression constructs; TH, structural modeling, quantum-chemical calculations, writing; SHH, study design, data analysis, quantum-chemical calculations, writing. All authors contributed to the editing of the manuscript.

## Data Availability Statement

The data that support the findings of this study are available from the corresponding author upon reasonable request.

## Expanded View Figures

**Figure EV1.**
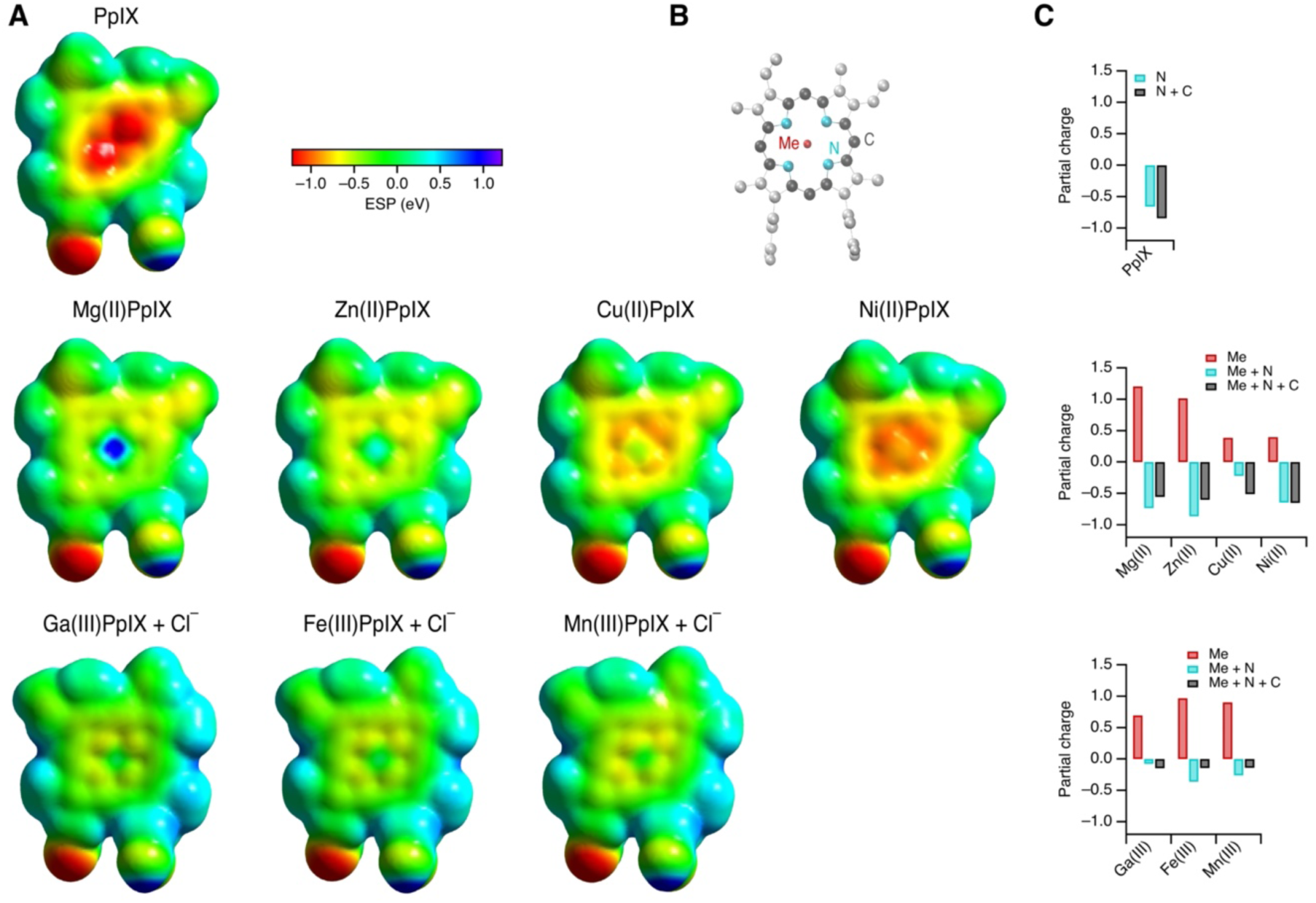
Electrostatic surface potential (ESP) of MePpIX. (**A**) Results of the DFT (density functional theory) calculations with Gaussian 16, using the aug-cc-pVDZ basis set (augmented double Dunning’s correlation consistent basis set) with water as the solvent. ESPs mapped onto electron density isosurfaces of 0.0004 atomic units are shown in electron volts (eV) according to the color scheme. The structures are grouped according to no metal center (PpIX, *top*), divalent ion configuration (*middle*), and trivalent ion configuration (*bottom*). In the latter cases, the structures also contain a chloride ion on the reverse side to compensate for the extra charge and to make the surface potentials comparable. In all cases, each of the carboxylate groups of the PpIX rings was saturated with an extra H^+^. The optimization was performed for Cu(II)PpIX and Zn(II)PpIX with default, for the remaining variants with a loose convergence criterion. (**B**) MgPpIX structure as an example to indicate which atoms were used to calculate the summed partial charges, as shown in (C). (**C**) Partial charges of the metal center (red), the metal center plus the surrounding ring of nitrogen atoms (cyan), and also including the carbon atoms of the PpIX ring (dark gray). ESP charges were determined from the Gaussian checkpoint files using MK (Merz-Singh-Kollman) population analysis and UFF (Universal Force Field) radii (Iop 6/41=10, 6/42=10).

**Figure EV2.**
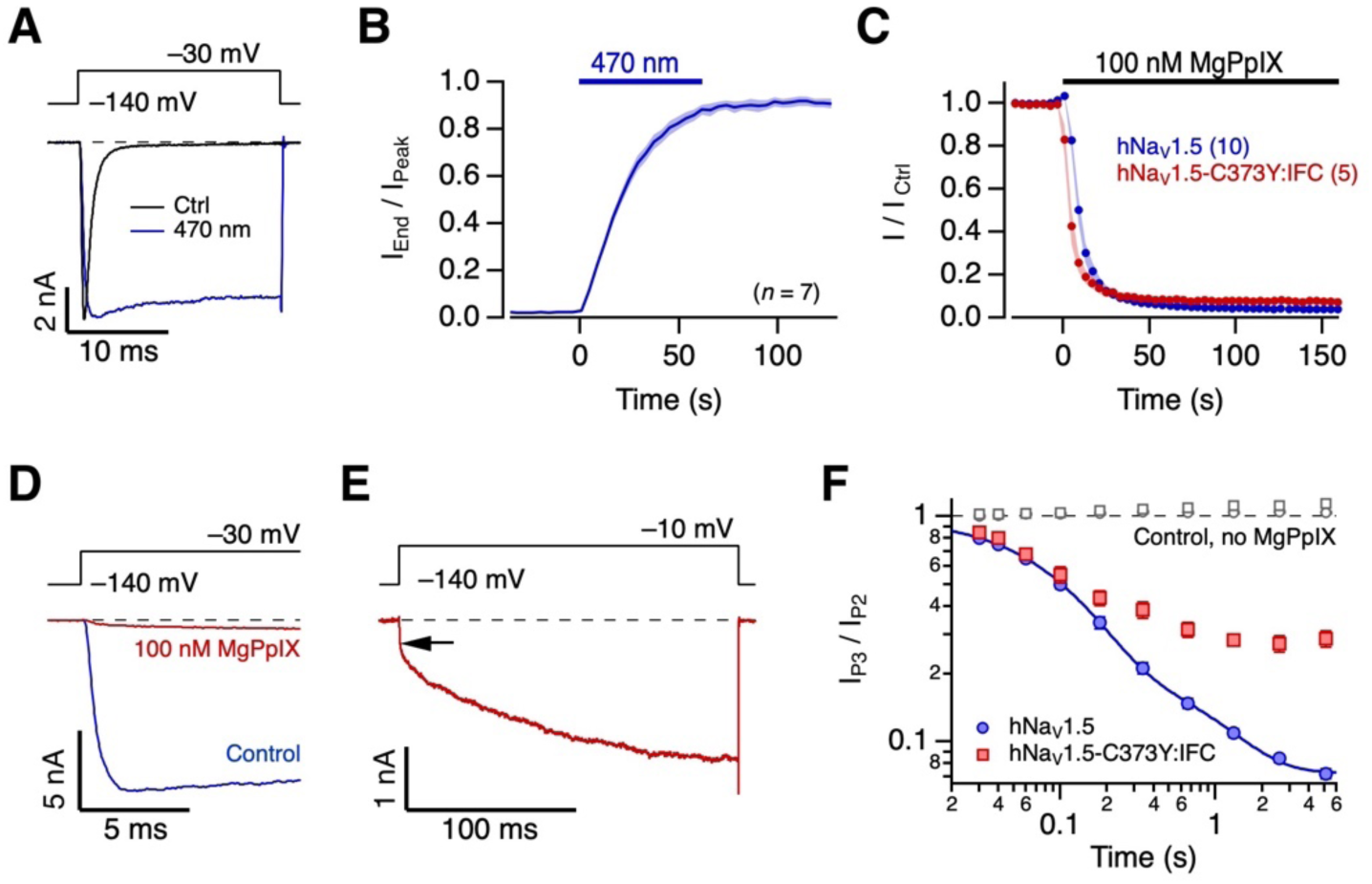
Light-inducible non-inactivating hNa_V_1.5 channels. Experiments were carried out with hNa_V_1.5-C373Y:IFC, i.e., a variant lacking a cysteine residue in the pore but possessing a cysteine instead of methionine in the inactivation domain “IFM” between domains III and IV, referred to as “IFC” to reflect the mutation M1487C. The variant was expressed in HEK293T cells, and currents were measured in the whole-cell mode. The pipette solution was supplemented with 250 µM lucifer yellow. (**A**) Representative current recording for the indicated step depolarization under control conditions (black) and after blue-light illumination of the cell filled with lucifer yellow (blue). (**B**) Mean time course of the current at the end of a 20-ms depolarization divided by the peak current (to serve as an index of inactivation loss) as a function of time. The horizontal bar indicates the illumination of the cell through a 20x objective with light from a 470-nm LED. The thick line represents the mean, and sem is indicated by shading. The loss of inactivation proceeded with a time constant of about 15 s. (**C**) Time course of normalized peak current at -30 mV with the application of 100 nM PpIX for hNa_V_1.5 (inactivating, blue) and hNa_V_1.5-C373Y:IFC channels after light-induced inactivation removal (red). Straight lines connect the data points for clarity. (**D**) Superposition of non-inactivating hNa_V_1.5-C373Y:IFC currents before (blue) and after application of 100 nM MgPpIX (red). (**E**) Current trace under the conditions shown in (D) for 200 ms visualizing the reversal of channel inhibition at -10 mV. The arrow approximately indicates the instantaneous current level before reverse use-dependence occurs. (**F**) Kinetics of channel block after partial relief from inhibition (as in Fig. 2E). Open symbols refer to control measurements in the absence of MgPpIX.

**Figure EV3.**
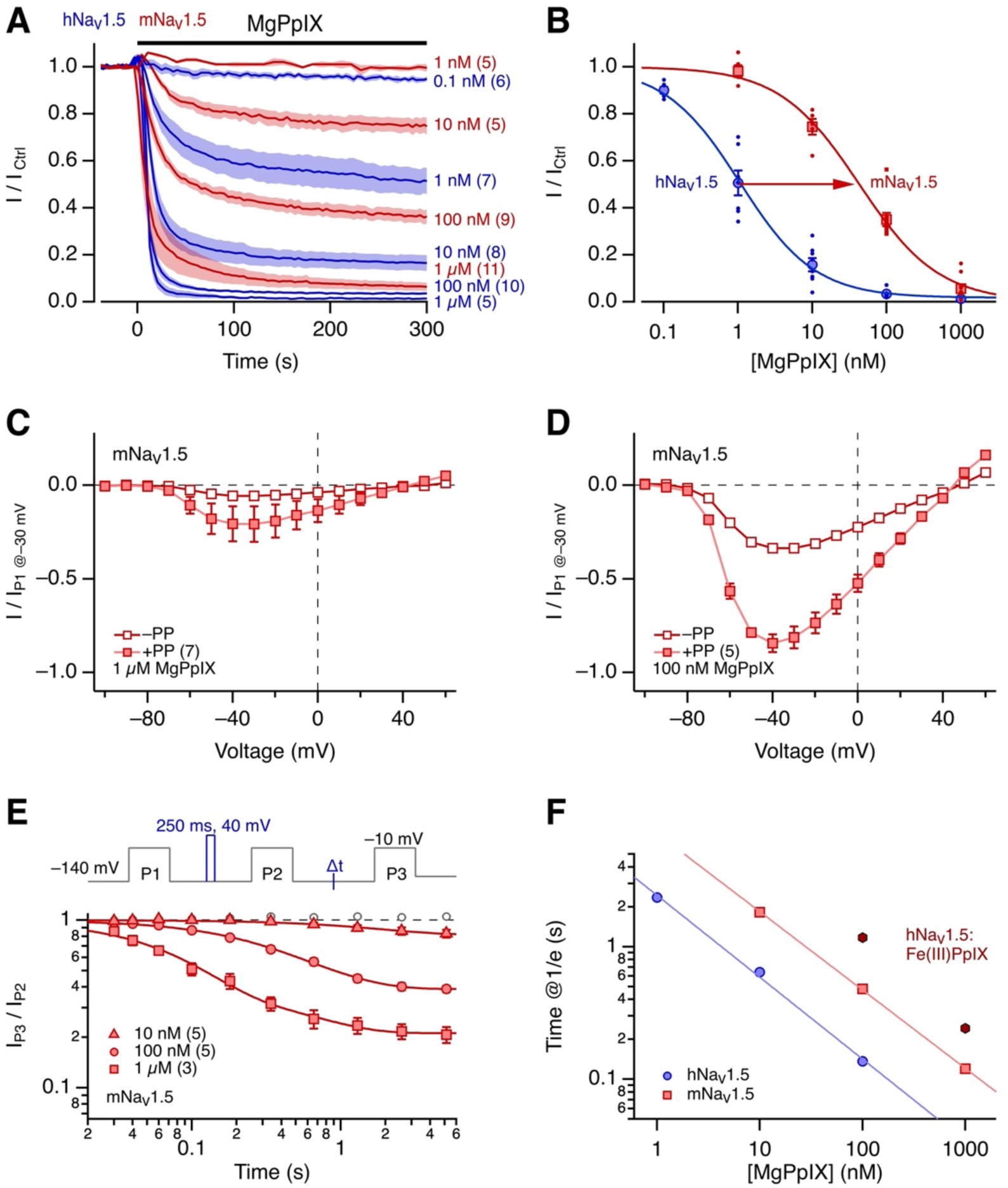
Inhibition of mouse Na_V_1.5 channels by MgPpIX. (**A**) Time course of peak current (at -30 mV) inhibition by the indicated concentrations of MgPpIX for human hNa_V_1.5 (blue, from Fig. 1) and mouse mNa_V_1.5 (red). Thick lines represent means, and sem is indicated by shading with *n* in parentheses. (**B**) Concentration-response of the normalized current after 300 s of MgPpIX application with superimposed Hill fits (Eq. 4). Data for hNa_V_1.5 are identical to those of Fig. 1; the results for mNa_V_1.5 are: IC_50_ = 42.8 ± 5.0 nM, *n*_H_ = 0.81 ± 0.07, and *a*_οο_ was constrained to 0. (**C**) Current–voltage relationships in the presence of 1 µM MgPpIX, normalized to the value obtained at -30 mV prior to MgPpIX application. Open symbols represent values of the first depolarizing pulse (P1), while filled symbols originate from the recordings following a prepulse to -10 mV for 100 ms (P2). Straight lines connect data points for clarity. (**D**) As in (C) but for 100 nM MgPpIX. (**E**) Kinetics of the onset of current inhibition after prepulse-induced reversal for mNa_V_1.5 at the indicated MgPpIX concentrations. Curves are the results of double-exponential fits. Open symbols refer to control conditions in the absence of MgPpIX. (**F**) From experiments as shown in (E) for mNa_V_1.5 and Fig. 2E for hNa_V_1.5, the time needed to inhibit the channels to 1/e (37%) is plotted as a function of MgPpIX concentration. Results for hNa_V_1.5 and hemin (Fe(III)PpIX) are also indicated. In this log-log presentation, the re-block kinetics are linear, with the characteristic time becoming about 4.3-times smaller at 10-fold concentration increase. For comparison, the reestablishment of the block was about 8 times slower for Fe(III)PpIX. A comparison of human and mouse Na_V_1.5 revealed a roughly 3.3-fold faster re-block by MgPpIX for the human isoform.

**Figure EV4.**
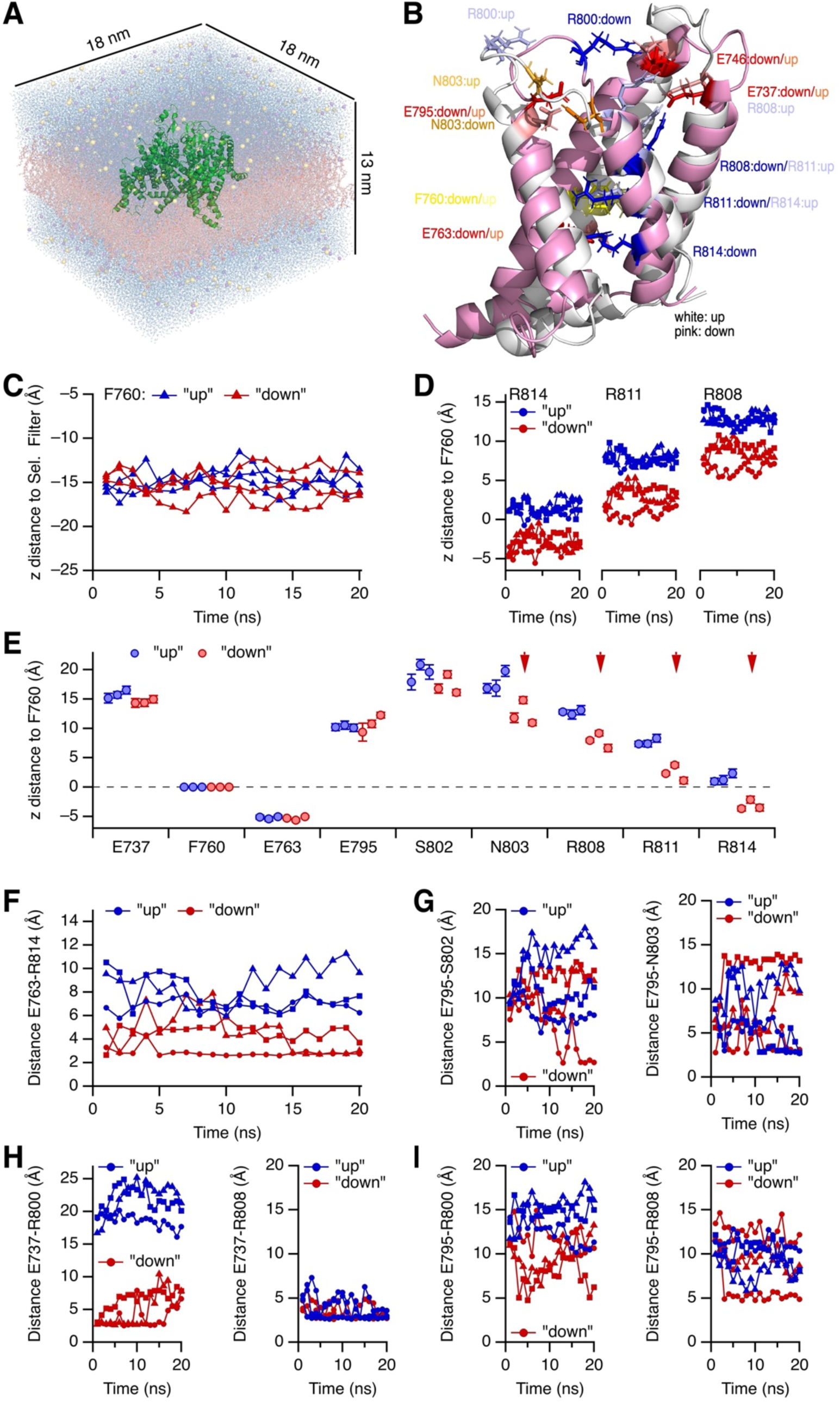
Structural model of hNa_V_1.5 with VSD II in a deactivated position. (**A**) Molecular dynamics simulation box containing the channel protein (green), a lipid bilayer (wheat), water molecules (blue), and ions (K^+^, purple; Cl^−^ yellow) with a total of about 470,000 atoms. Periodic boundary conditions were applied during simulation. (**B**) Superposition of VSD II in “up” (white) and “down” (pink) configuration. The “up” configuration is the structure as obtained from the cryo-EM data (PDB: 6LQA), the deactivated “down” configuration was obtained by aligning VSD II to the voltage sensor of Kv7.1 channels with subsequent energy minimization. For the superposition, structures were aligned according to the S6 segments (CA atoms only) of all four domains. (**C**-**I**) Distance measurements of various residues from each three MD simulations of 20 ns in the “up” and “down” configuration of VSD II. (**C**) z distance of the β carbon (CB) of F760 relative to the mean CB position of the selectivity filter “DEKA” (D372, E901, K1419, A1711) from all four domains, indicating the relative stability of the F760 position within the channel protein. (**D**) z distance of the CB carbons of R814, R811, and R808 relative to CB of F760. (**E**) Average values (means of the structures from 10-20 ns, ±s.d.) of the z distances of the indicated residues (always the CB carbon) to F760/CB for three simulation systems in the “up” and three in the “down” configuration. Minimum distance of the amide groups of R814 (NH1, NH2) to the oxygens of E763 (OE1, OE2); R814 and E763 form an ion–ion bond in the “down” configuration. (**G**) Distances of E795 to S802 (*left*) and N803 (*right*). (**H**) Distances of E737 to R800 (*left*) and R808 (*right*). (**I**) Distances of E795 to R800 (*left*) and R808 (*right*).

**Figure EV5.**
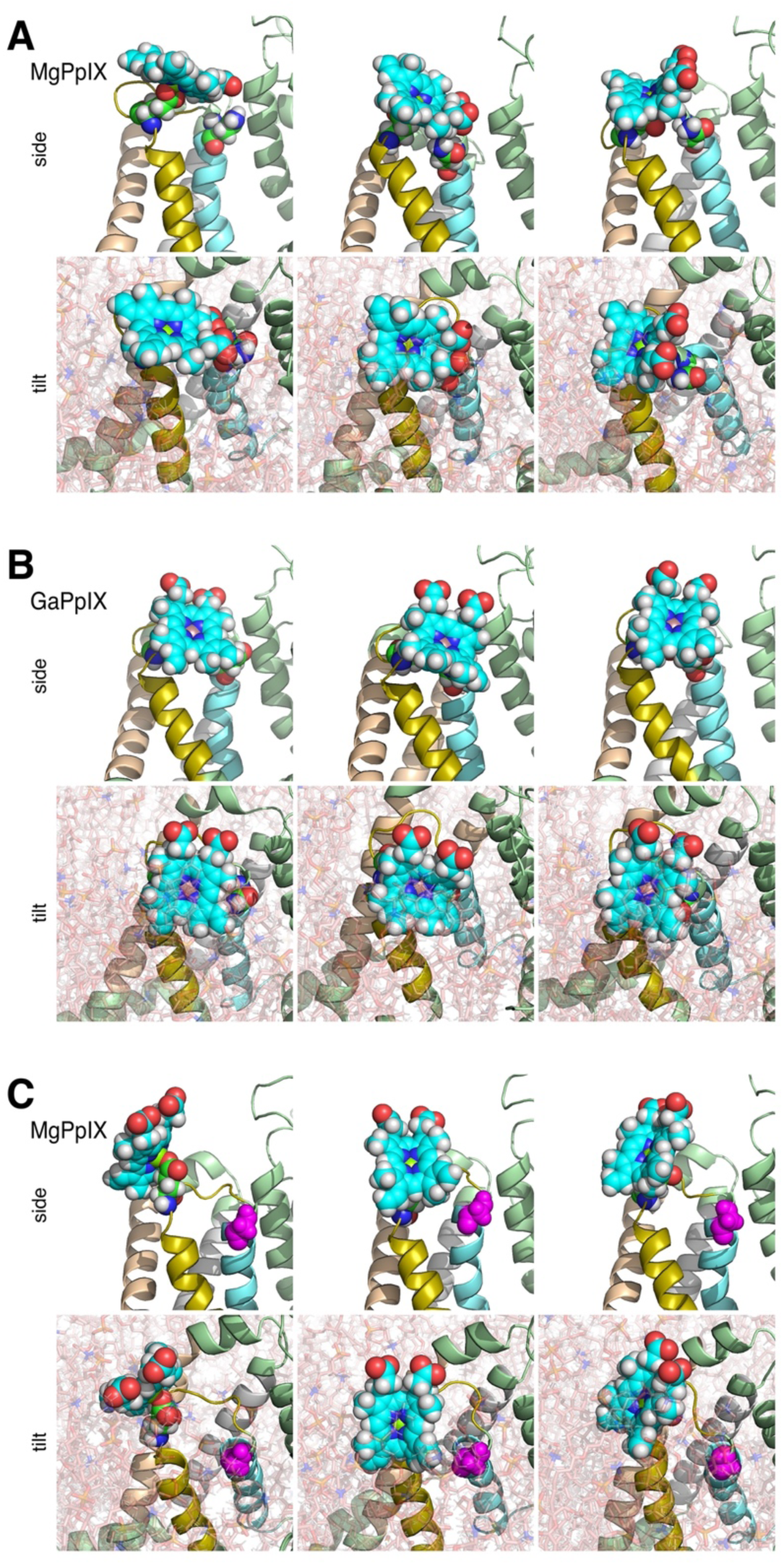
Possible binding configuration of MePpIX at the voltage sensor of domain II. (**A-C**) Three snapshots, one from each of the three molecular dynamics simulation runs, as detailed in Fig. EV4 with MgPpIX (A, C) or GaPpIX (B) placed near VSD II. Channel structures are hNa_V_1.5 with VSD II in a deactivated “down” position (A, B) and the same for channel variant N803G (C). *top*, Presentations of VSD II in side view without water, ions, and lipids. *bottom*, The same structures as in the *top* panels but including lipids, and tilted by about 45°. Residues E795 and N803 are shown as spheres with carbon atoms in green. G803 (in C) is shown in magenta. The carbon atoms of the MePpIX are shown in cyan.

