## Supplemental Information for "Voltage-sensor trapping of cardiac Na^+^ channels by Mg-protoporphyrin impairs cancer cell migration"

**Supplementary Material**

Mahdi Jamili[a], Marwa Ahmed[a], Alisa Bernert[a], Johann Rößler[a], Guido Gessner[a], Roland Schönherr[a], Toshinori Hoshi[b], and Stefan H. Heinemann[a,*]

[a] *Center for Molecular Biomedicine, Department of Biophysics, Friedrich Schiller University Jena and Jena University Hospital, Jena, Germany*

[b] *Department of Physiology, University of Pennsylvania, Philadelphia, PA 19104-6085, USA*

* Corresponding author
*Center for Molecular Biomedicine, Department of Biophysics, Friedrich Schiller University Jena and Jena University Hospital, Hans-Knöll-Straße 2, 07745 Jena, Germany*

ORCID: 0000-0002-4144-0251

### Supplementary Tables

**Supplementary Table 1.** Parameters describing the voltage dependence of activation and inactivation, as well as the remaining current at -30 mV (*Rem. I*) after the application of 100 nM MgPpIX for $Na_V$ channels and variants.

| **Channel type** | ***V*m (mV)** | ***k*m (mV)** | ***n*** | ***V*h (mV)** | ***k*h (mV)** | ***n*** | **Rem. I in 100 nM MgPpIX (%)** | ***n*** |
|---|---|---|---|---|---|---|---|---|
| $hNa_V1.5$ | -55.6 ± 0.6 | 8.8 ± 0.2 | 39 | -94.2 ± 0.8 | 5.7 ± 0.1 | 40 | 3.4 ± 0.4 | 10 |
| $nhNa_V1.5$ | -48.9 ± 2.5 | 9.8 ± 0.7 | 6 | -90.3 ± 1.8 | 5.7 ± 0.2 | 6 | 3.8 ± 0.7 | 5 |
| $mNa_V1.5$ | -59.5 ± 0.9 | 8.7 ± 0.2 | 19 | -94.7 ± 0.9 | 5.6 ± 0.1 | 21 | 35.1 ± 2.8 | 9 |
| $hNa_V1.2$ | -40.1 ± 0.9 | 8.0 ± 0.5 | 9 | -70.1 ± 1.5 | 6.4 ± 0.6 | 9 | 105.7 ± 2.0 | 5 |
| $hNa_V1.4$ | -40.8 ± 0.6 | 8.3 ± 0.4 | 9 | -76.5 ± 0.6 | 5.6 ± 0.2 | 10 | 101.8 ± 3.3 | 5 |
| $hNa_V1.7$ | -40.5 ± 1.2 | 8.6 ± 0.3 | 11 | -85.5 ± 1.1 | 5.8 ± 0.2 | 11 | 106.5 ± 3.7 | 6 |
| $hNa_V1.5$ variants: | | | | | | | | |
| C373Y | -54.1 ± 1.4 | 8.0 ± 0.4 | 5 | -92.9 ± 1.4 | 5.5 ± 0.2 | 5 | 5.4 ± 1.2 | 5 |
| E737A | -45.1 ± 1.9 | 9.4 ± 0.3 | 5 | -94.6 ± 1.8 | 5.4 ± 0.2 | 5 | 13.3 ± 2.5 | 5 |
| N740P | -57.0 ± 1.5 | 8.8 ± 0.5 | 10 | -96.2 ± 1.8 | 6.0 ± 0.2 | 11 | 5.6 ± 0.6 | 5 |
| N740Q | -53.3 ± 1.2 | 8.6 ± 0.4 | 5 | -91.1 ± 0.9 | 5.7 ± 0.3 | 5 | 6.5 ± 0.7 | 5 |
| E743A | -57.7 ± 0.9 | 8.4 ± 0.5 | 6 | -98.2 ± 1.6 | 5.5 ± 0.2 | 7 | 5.1 ± 1.1 | 7 |
| S743E | -55.4 ± 1.6 | 8.5 ± 0.3 | 5 | -95.1 ± 1.7 | 6.0 ± 0.4 | 5 | 32.4 ± 3.3 | 5 |
| E746S | -49.3 ± 2.2 | 10.0 ± 0.8 | 5 | -91.0 ± 1.7 | 5.3 ± 3.1 | 5 | 4.1 ± 1.2 | 5 |
| E746K | -59.8 ± 1.5 | 8.6 ± 0.5 | 6 | -100.3 ± 1.4 | 5.4 ± 0.2 | 7 | 7.4 ± 0.9 | 7 |
| E795A | -48.0 ± 1.2 | 12.9 ± 0.3 | 8 | -96.9 ± 1.8 | 5.7 ± 0.2 | 8 | 100.0 ± 2.3 | 5 |
| R800A | -55.6 ± 2.4 | 10.3 ± 0.8 | 5 | -100.4 ± 3.2 | 5.6 ± 3.2 | 5 | 6.8 ± 1.8 | 5 |
| R800D | -63.2 ± 2.2 | 8.9 ± 0.4 | 5 | -103.0 ± 1.7 | 5.5 ± 0.4 | 5 | 15.2 ± 3.3 | 6 |
| S802G | -54.2 ± 2.1 | 9.4 ± 0.8 | 5 | -97.1 ± 1.7 | 6.0 ± 0.4 | 5 | 27.7 ± 4.1 | 5 |
| S802E | -64.7 ± 1.3 | 9.3 ± 9.1 | 5 | -108.5 ± 3.0 | 6.4 ± 0.4 | 5 | 26.7 ± 2.6 | 5 |
| N803G | -51.9 ± 0.9 | 9.6 ± 0.5 | 5 | -95.4 ± 0.9 | 5.5 ± 0.1 | 5 | 93.8 ± 1.2 | 5 |
| N803A | -51.9 ± 1.8 | 9.6 ± 0.2 | 5 | -98.3 ± 1.3 | 6.1 ± 0.5 | 5 | 90.3 ± 4.6 | 6 |
| N803Q | -56.2 ± 1.1 | 8.1 ± 0.3 | 5 | -94.4 ± 2.0 | 5.5 ± 0.3 | 5 | 5.5 ± 2.1 | 5 |
| N803S | -55.9 ± 1.0 | 9.5 ± 0.5 | 8 | -98.0 ± 1.7 | 5.9 ± 0.2 | 8 | 89.0 ± 2.3 | 6 |
| N803F | -42.3 ± 1.2 | 13.8 ± 0.2 | 8 | -98.9 ± 1.3 | 5.6 ± 0.2 | 8 | 80.8 ± 2.9 | 7 |
| N803H | -54.1 ± 1.7 | 10.4 ± 0.3 | 5 | -98.3 ± 2.9 | 6.4 ± 0.5 | 5 | 6.5 ± 1.0 | 8 |
| N803D | -60.9 ± 0.6 | 7.7 ± 0.6 | 7 | -98.4 ± 1.9 | 5.6 ± 0.3 | 7 | 94.5 ± 2.0 | 5 |
| N803R | -54.2 ± 2.0 | 9.8 ± 0.6 | 5 | -101.2 ± 2.6 | 6.2 ± 0.2 | 5 | 12.2 ± 1.5 | 5 |
| N803K | -50.6 ± 0.9 | 9.5 ± 0.2 | 5 | -94.6 ± 3.0 | 6.8 ± 0.5 | 5 | 18.5 ± 1.9 | 5 |
| S802E:N803G | -60.1 ± 1.7 | 7.9 ± 0.6 | 5 | -100.3 ± 1.3 | 6.0 ± 0.4 | 5 | 90.1 ± 2.4 | 5 |
| R808A | -44.8 ± 1.1 | 11.8 ± 0.3 | 5 | -100.0 ± 0.8 | 6.8 ± 0.5 | 5 | 58.5 ± 5.5 | 5 |
| $hNa_V1.7$ variants: | | | | | | | | |
| E829S | -38.2 ± 0.3 | 10.6 ± 0.2 | 5 | -90.7 ± 1.7 | 6.1 ± 0.3 | 5 | 97.3 ± 10.4 | 5 |
| G830N | -40.3 ± 2.2 | 9.9 ± 0.6 | 6 | -90.1 ± 1.7 | 5.9 ± 0.4 | 6 | 106.1 ± 5.1 | 6 |
| E829S:G830N | -58.5 ± 1.1 | 9.2 ± 0.4 | 10 | -98.5 ± 0.7 | 5.8 ± 0.3 | 10 | 7.5 ± 1.4 | 5 |

**Supplementary Figures**

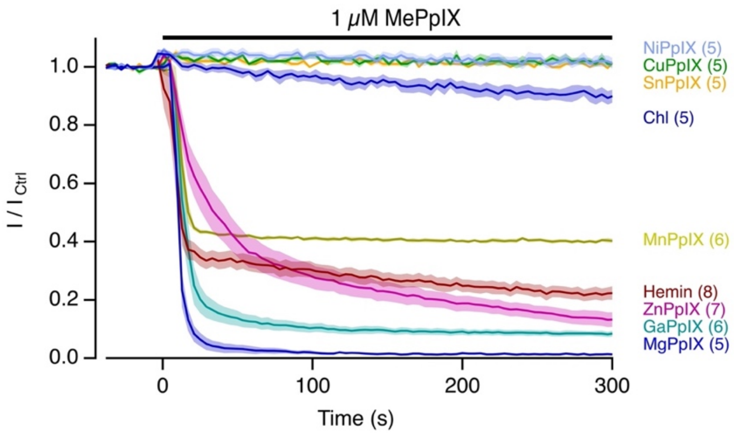


**Supplementary Figure 1. Kinetics of hNa$_V$1.5 inhibition by MePpIX.**

The time course of the normalized peak current at -30 mV with the application of the indicated metal protoporphyrins (MePpIX) or chlorophyll-A (Chl) at 1 µM. The thick lines denote mean values, and sem is indicated by shading, with *n* in parentheses.

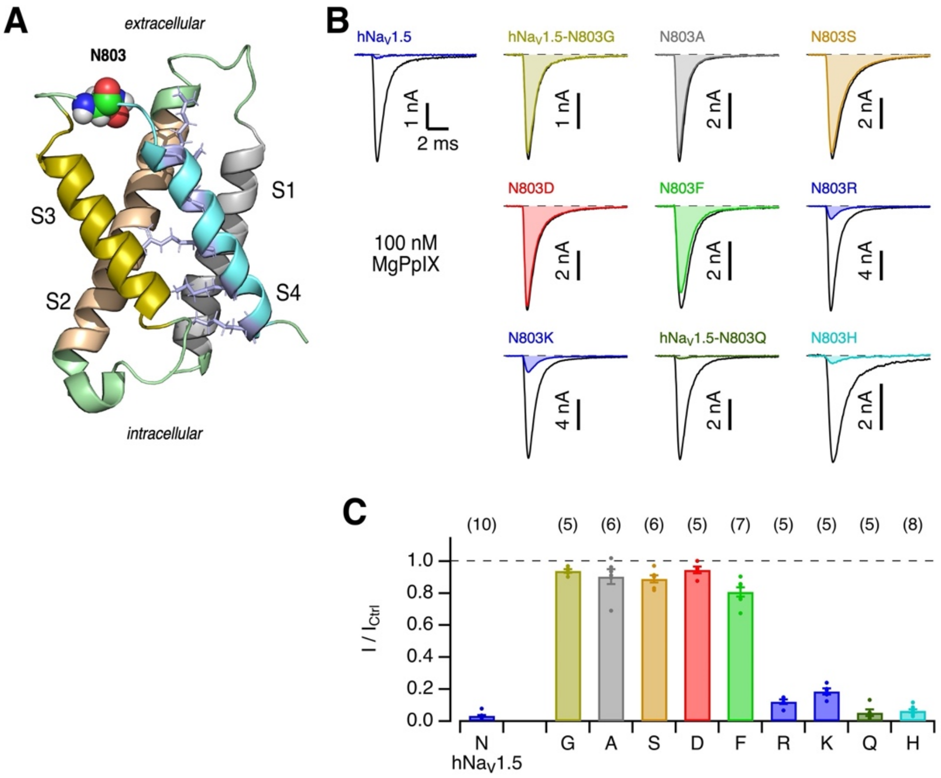


**Supplementary Figure 2.** **Mutagenesis of hNa$_V$1.5 at site N803.**

(**A**) Structure of the hNa$_V$1.5 domain-II voltage sensor (from PDB: 6LQA) with residue N803 highlighted: carbon, green; nitrogen, blue; oxygen, red; hydrogen, grey. Positively charged residues of S4 are shown as sticks (light blue). (**B**) Representative current recordings at -30 mV for the indicated channel types and variants before (black) and 5 min after the application of 100 nM MgPpIX (colored). (**C**) Mean fractional peak current at -30 mV remaining 5 min after the application of 100 nM MgPpIX for wild type hNa$_V$1.5 and variants mutated at site N803. Data are means ± sem (*n* in parentheses, individual results indicated as dots).

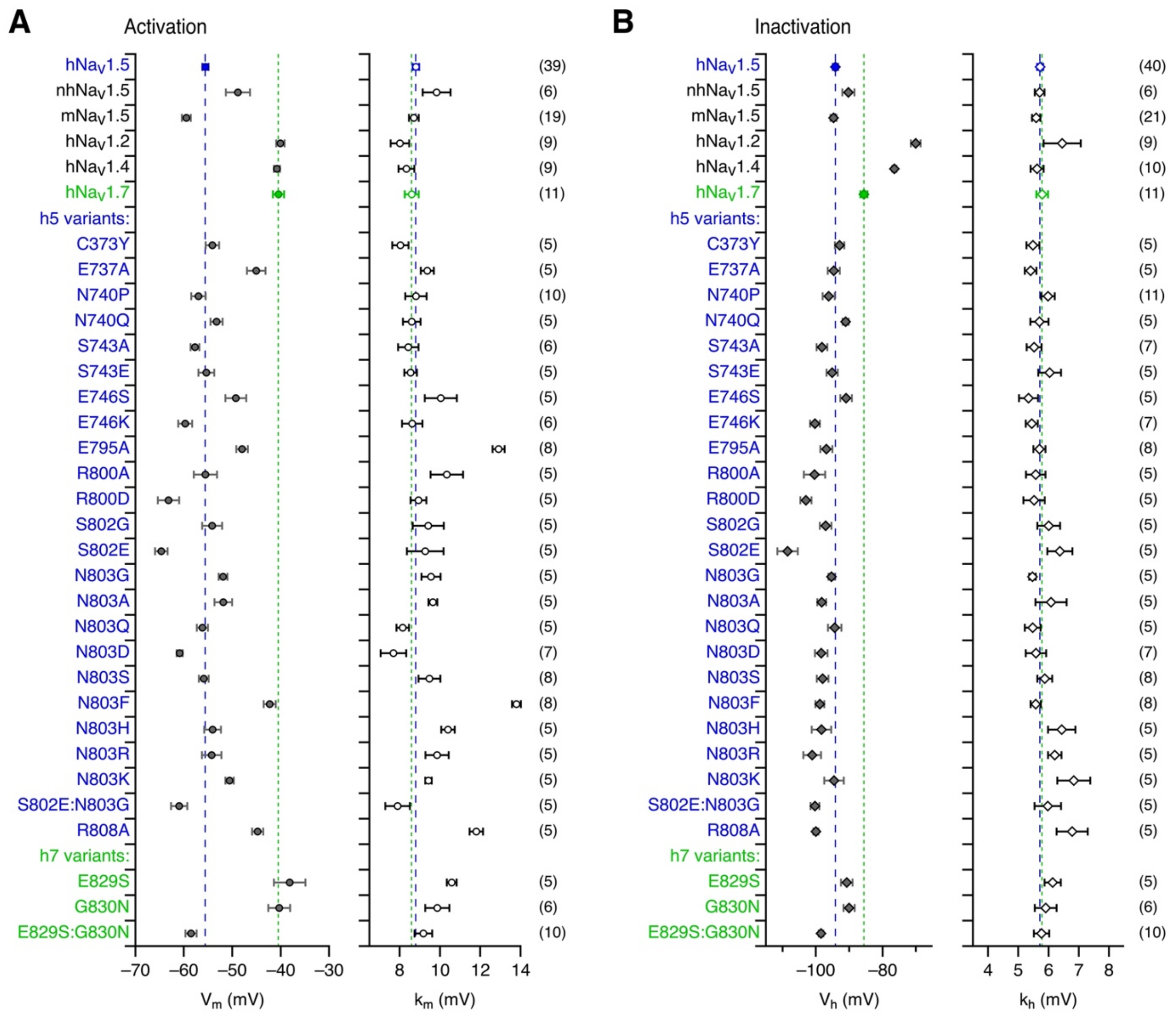


**Supplementary Figure 3. Parameters describing the voltage dependence of activation and inactivation of $Na_V$ channels and their variants.**

(**A**) $V_m$ and $k_m$ as results of the analysis of current–voltage relationships (Eq. 1) for the indicated $Na_V$ channel types and variants. (**B**) $V_h$ and $k_h$ values according to Eq. 2 describing the voltage dependence of channel inactivation caused by 500-ms prepulses. Data are means ± sem with *n* in parentheses. Most notably, mutation S802E caused a marked left-shift in the voltage dependence of activation and inactivation in $hNa_V1.5$. Moreover, only the combination of the mutations E829S and G830N in $hNa_V1.7$, but not the individual mutations, altered the half-maximal activation voltage ($V_m$) from -40.5 ± 0.1 mV (*n* = 11) for $hNa_V1.7$ to -58.5 ± 0.1 mV (*n* = 10), which is close to the value obtained for $hNa_V1.5$ (-55.6 ± 0.1 mV, *n* = 39).

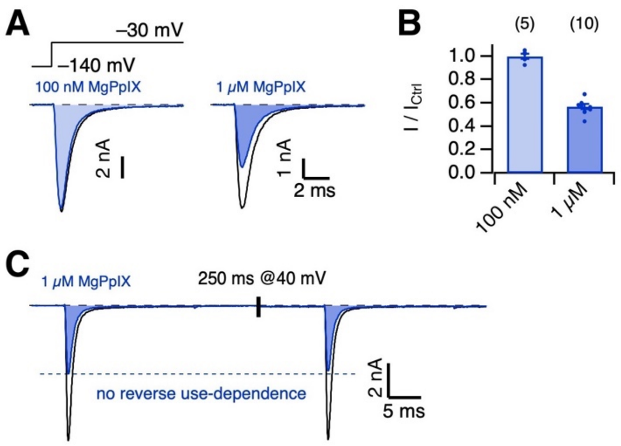


**Supplementary Figure 4. hNa$_V$1-5-E795A is only weakly inhibited by MgPpIX.**

(**A**) Pulse protocol and superimposed current traces of hNa$_V$1.5-E795A before (black) and after application of the indicated concentration of MgPpIX (blue). (**B**) Mean remaining current at -30 mV after application of MgPpIX. Data are means ± sem (*n* in parentheses), with individual data points indicated as dots. (**C**) Current response of consecutive pulses to -30 mV with a depolarizing episode (250 ms at 40 mV) between the pulses, indicated by the vertical bar. Black traces were recorded under control conditions, blue traces after application of 1 μM MgPpIX.

**Supplementary Movies**

1) Three MD simulations of $hNa_V1.5$ with VSD II in a “down” position and MgPpIX. E795 is shown as spheres with atom colors, N803 in orange.
2) Three MD simulations of $hNa_V1.5$ with VSD II in a “down” position and GaPpIX. E795 is shown as spheres with atom colors, N803 in orange.
3) Three MD simulations of $hNa_V1.5$ with VSD II in a “down” position and NiPpIX. E795 is shown as spheres with atom colors, N803 in orange.
4) Three MD simulations of $hNa_V1.5$-N803G with VSD II in a “down” position and MgPpIX. E795 is shown as spheres with atom colors, G803 in orange.
5) Three MD simulations of $hNa_V1.5$-E795A with VSD II in a “down” position and MgPpIX. A795 is shown as spheres with atom colors, N803 in orange.

(All files are integrated in one pptx document.)
